# Cariogenic *Streptococcus mutans* produces strain-specific antibiotics that impair commensal colonization

**DOI:** 10.1101/755025

**Authors:** Xiaoyu Tang, Yuta Kudo, Jonathon Baker, Sandra LaBonte, Peter A. Jordan, Shaun M. K. McKinnie, Jian Guo, Tao Huan, Bradley S. Moore, Anna Edlund

## Abstract

*Streptococcus mutans* is a common constituent of dental plaque and an etiologic agent of dental caries (tooth decay). Here we elucidate a biosynthetic pathway, encoded by globally distributed strains of *S. mutans*, which produces a series of bioactive small molecules including reutericyclin and two *N*-acyl tetramic acid analogues active against oral commensal bacteria. This pathway may provide *S. mutans* with a competitive advantage, promoting dysbiosis and caries pathogenesis.

---

The human microbiota consists of trillions of symbiotic microbial cells that not only help its host to digest dietary components(1-3), metabolize drugs(4-6), and regulate immune system(7-9) but also produce complex small molecules such as antimicrobial nonribosomal peptides (NRPs) and polyketides (PKs)(10-12). Recent computational mining efforts of genomes and metagenomes of the human microbiome revealed a vast diversity (∼14,000) of putative biosynthetic gene clusters (BGCs) encoding small molecules across all human body sites(13), of which many represented NRP, PK and hybrid NRP-PK small molecules. A typical gut harbors 599 BGCs, while a typical oral cavity harbors 1061 BGC(13). Thus far, most research efforts have focused on characterization of BGCs and small molecules from the gut microbiome, leaving large knowledge gaps of crucial signaling molecules of the oral cavity.

The oral cavity harbors a high species diversity with over 700 bacterial species, which mainly colonize four physically distinct niches including dental plaque, tongue dorsum, buccal mucosa, and saliva(14). Residents of the dental plaque have been implicated in a variety of diseases, including dental caries, which affects more than a third of the world’s population and results in approximately $300 billion in direct treatment costs to the global economy annually(15-17). Although caries is a polymicrobial disease caused by a dysbiosis in the dental plaque microbial community, *Streptococcus mutans*, with its copious acid production and prodigious biofilm formation, is still considered a primary etiologic agent(18, 19). To persist in the dental plaque community and cause disease, *S. mutans* must be able to outcompete commensal bacteria directly.

Small molecules produced by BGCs are increasingly recognized to play major roles in species-species communication and interactions(10, 13), and a recent study predicted 355 strain-specific BGCs across 169 *S. mutans* genomes(20). Although the production of mutactins (a group of bacteriocins) has been recognized for contribution to the colonization and establishment of *S. mutans* in the dental biofilm(21), the roles of other genetically encoded small molecules in *S. mutans* is barely explored, with exception of the mutanobactins. Mutanobactins are compounds of hybrid polyketide synthase and nonribosomal peptide synthetase (PKS/NRPS) origin that inhibit the morphological transition of *Candida albicans*(22). Previous bioinformatics efforts identified an orphan hybrid PKS/NRPS BGC (recently designated *muc*)(20, 23), that is distributed among a subset of *S. mutans* strains. Within *muc*, five biosynthetic proteins are highly homologous (48%-69%) to cognates involved in the biosynthesis of reutericyclin (RTC)(24) (**Supplementary Fig. 1** and **Fig. 1a**). RTC, which originated from the sourdough isolates of *Lactobacillus reuteri*, acts as a proton ionophore antibiotic that modulates the microbial community of sourdough(25, 26). Interestingly, we found that *S. mutans* strains encoding *muc* were dispersed geographically and frequently associated with severe dental caries (**Supplementary Table 1**). The goal of this study was to determine whether *muc* produces RTC or RTC-like molecules, and if these molecules can affect inter-species competition in the oral cavity.

**Figure 1.**
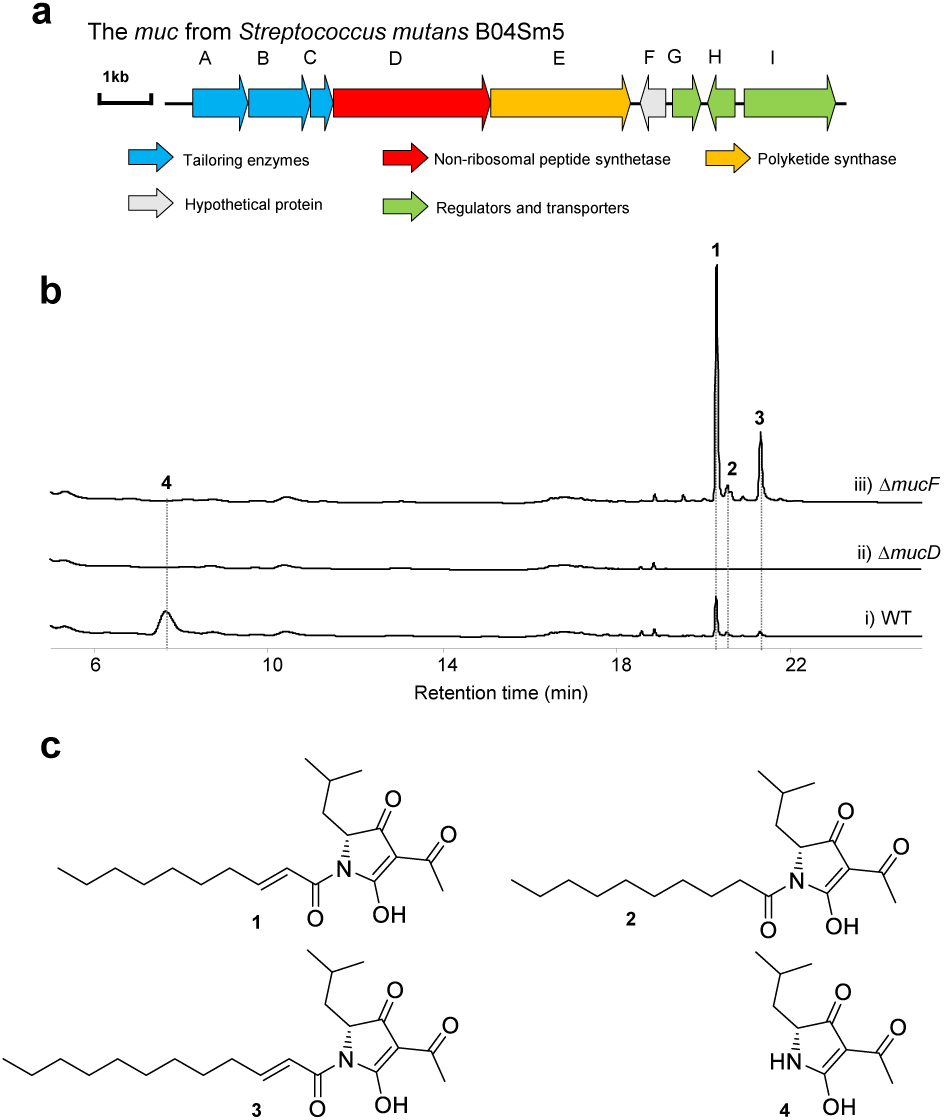
Identification of an orphan gene cluster from *S. mutans* and its metabolites. (**a**) Mutanocyclin gene cluster (*muc*) annotation. (**b**) HPLC profile of extracts from wild type (WT) *S. mutans* B04Sm5 (i), *S. mutans* B04Sm5*/*Δ*mucD* (ii), and *S. mutans* B04Sm5*/*Δ*mucF* (iii). (**c**) Structures of metabolites identified from *S. mutans* in this study, including reutericyclin A (**1**), reutericyclin B (**2**), reutericyclin C (**3**), and mutanocyclin (**4**). C, condensation; A, adenylation; T, thiolation; KS, ketosynthase; TE, thioesterase.

To determine the product of *muc*, we utilized homologous recombination to delete the gene *mucD*, which encodes an assembly line NRPS tridomain protein (**Fig. 1a**). We conducted all our biosynthetic experiments in *S. mutans* B04Sm5, a strain bearing *muc*, which was isolated from a child with severe early childhood caries(27). The wild type (WT) strain and Δ*mucD* mutant were cultured and extracted for HPLC analyses. The results showed that the WT strain produced four metabolites not present in the Δ*mucD* mutant (**Fig. 1b**). These molecules were purified via preparative HPLC (**Supplementary Note 1**) and characterized by High-Resolution Mass Spectrometry (HR-MS/MS), chemical synthesis, and/or one- and two-dimensional Nuclear Magnetic Resonance (NMR) experiments to yield a group of tetramic acids, including RTC (renamed RTC A, **1**), two new RTC analogues (RTC B (**2**) and RTC C (**3**)), and a new organic acid (**4**) (**Fig. 1c, Supplementary Note 1, Supplementary Tables 2-3, Supplementary Figs. 2-22**). During preparation of this manuscript, Chen and colleagues published the identification of **4** (designated mutanocyclin (MUC)) using a new heterologous genetic system(23); however, production of **1**-**3** in both heterologous host and wild type producers was not reported.

*Muc* is a ∼13 kb hybrid NRPS-PKS pathway encoding nine proteins (**Supplementary Table 5**). *In silico* analysis revealed that *mucD* to *mucE* encode the core assembly line protein machinery (**Fig. 1a** and **Fig. 2a**). While MucD is a C-A-T tridomain protein, with specificity for adenylating leucine, MucE contains an KS-T-TE module, as commonly present in the termination modules of PKS assembly lines. Based on the enzymatic logic of thiotemplate-mediated assembly line biosynthesis, we propose that **1**-**3** are assembled, respectively, from *trans*-2-decenoyl-ACP, decenoyl-ACP, and *trans-*2-dodecenoyl-ACP starter units through elongation with leucine, followed by elongation with a malonyl-CoA extender unit (**Fig. 2a**). Inspection of structures of **1**-**3** suggested that the A domain of MucD appears to install a *D*-leucine residue into the final product. To explore this hypothesis, we fed both [^13^C_1_] *L*- and *D*-leucine to cultures of *S. mutans* B04Sm5. MS analyses of the purified **1** and **3** only revealed the incorporation of [^13^C_1_] *L*-leucine (**Supplementary Figs. 23-24**). The same result was observed by feeding the original RTC producer *L. reuteri* with the same isomers (**Supplementary Fig. 25**). These results indicate that an unrecognized epimerization reaction is involved in **1**-**3** biosynthesis; however, no standard epimerization (E) domain or dual functioning C/E domains could be found either in the assembly line or encoded elsewhere in the BGC. Additionally, although a dual-function TE/E domain has been characterized from the nocardicin (NOC) biosynthetic assembly line(28), MucTE shows very low homology (20%/35%, identity/similarity) to the dual functioning NocTE domain.

**Figure 2.**
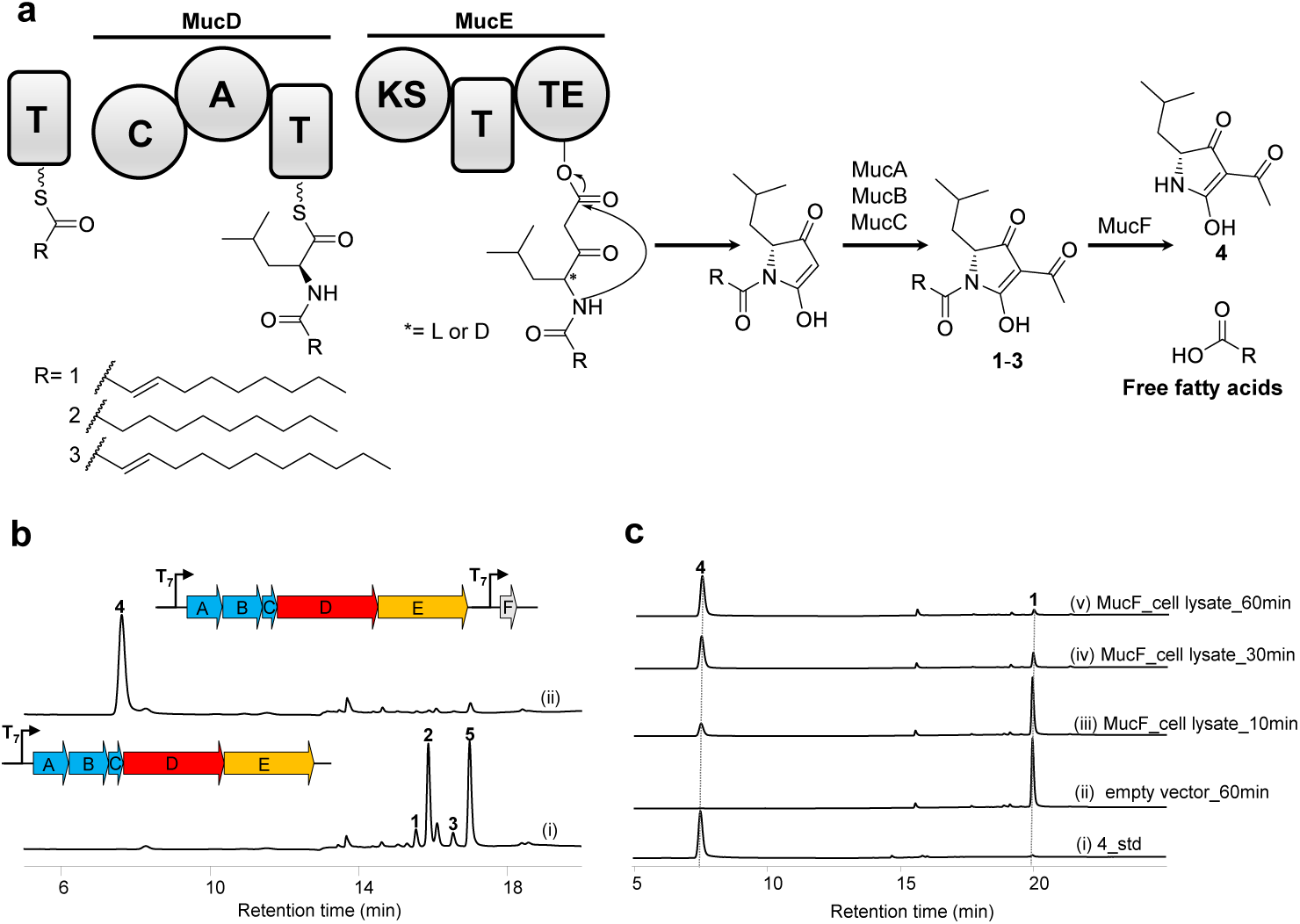
Model for 1-4 biosynthesis and characterization of MucF as deacetylase. (**a**) Model for the biosynthesis of **1**-**4**. (**b**) HPLC profiles of extracts from *E. coli* BAP1/pEXT06 (*mucA-D*) (i) and *E. coli* BAP1/pEXT07 (*mucA-D*+*mucF*) (ii). (**c**) HPLC analysis of (i) isolated **4** as a standard, (ii) compound **1** incubated with *E. coli* Rosetta2™(DE3)pLys/pET28a (empty vector) cell lysate for 60 min, compound **1** incubated with *E. coli* Rosetta2™(DE3)pLys/pEXT26 (carrying *mucF*) for 10 min (i), 30 min (ii), and 60 min (iii).

The first three genes (*mucA-C*) encode a hydroxymethylglutaryl-CoA synthase (MucA), a thiolase (MucB), and a hypothetical protein (MucC) (**Supplementary Table 5**), which show homology to the three submits (PhlA, PhlB and PhlC, respectively) of a multicomponent C-acetyltransferase involved in the acetylation of the type III PKS product phloroglucinol from *Pseudomonas fluorescens* Q2-87(29, 30). As the combination of the three genes was also identified in *rtc* (*rtcA, rtcC* and *rtcB*) from *L. reuteri* (24), the function of MucA–C is consistent with introducing the acetyl group to the pyrrolidine ring of **1**-**3** (**Fig. 2a**). We additionally annotated four genes downstream of the *mucA-E* operon encoding a small HXXEE domain-containing membrane-protein (MucF) of unknown function, two TetR/AcrR family transcriptional regulators (MucG and MucH), and one multidrug efflux pump (MucI) (**Supplementary Table 5**). Presumably, they are not involved in the direct synthesis of **1**-**3**. To verify this hypothesis, we cloned the operon from *mucA* to *mucE* into the pACYCDuet-1 vector to generate the plasmid pEXT06, in which the operon is exclusively under control of a T_7_ promoter. As expected, expression of *mucA-E* in *Escherichia coli* BAP1 strain resulted in the production of at least four products, including **1**-**3** and new compound **5** (**Fig. 2b** and **Supplementary Fig. 26**). **5** was purified via preparative HPLC and its structure was further confirmed as a new RTC analogue (RTC D) possessing a saturated C-12 fatty acid side chain by MS and NMR analyses (**Supplementary Table 4, Supplementary Fig. 26** and **Supplementary Figs. 27-30**). This result indicates that the first six genes *mucA-E* indeed compose the minimal BGC for **1**–**3** production (**Fig. 2a**).

As the structure of **4** is consistent with the RTC core lacking the fatty acyl chain, we first set out to detect whether the free fatty acid, *trans*-2-decenoic acid per compound **1**, is present in the extract of *S. mutans* B04Sm5. HPLC analyses confirmed that *S. mutans* B04Sm5 readily produced *trans*-2-decenoic acid (**Supplementary Fig. 31**). In contrast, it was not detected in the pathway-deficient mutant *S. mutans* B04Sm5/Δ*mucD*. These findings suggested that **4** may be derived from **1**-**3** via deacylation by an unknown enzyme. Interestingly, *trans*-2-decenoic acid is a known *Streptococcus* diffusible signal factor (SDSF) isolated from many *Streptococcus* species(31), which inhibits the hyphal formation of the opportunistic fungus *Candida albicans*. Among the annotated pathway enzymes, only the function of MucF was unassigned. The MucF protein sequence was subjected to a secondary structure prediction-based homology search (Phyre2), which suggested it is a polytopic (five) transmembrane α-helical protein (**Supplementary Fig. 32**) with low similarity (15%) to a viral protein (PDB 3LD1) with putative hydrolase activity. To explore whether MucF might be involved in the deacylation of **1**-**3**, we generated a *mucF* deletion mutant in *S. mutans* B04Sm5. HPLC analysis of the extract of mutant cultures showed that the Δ*mucF* mutant not only increased the production of **1**-**3** by ∼3-5 fold, but also lost the ability to produce **4** (**Fig. 1b**). These findings strongly suggested that MucF is essential for converting **1**-**3** to **4**. To further evaluate the function of MucF, we cloned and expressed *mucF* in *E. coli* and incubated **1** with the *E. coli*/*mucF* cell lysate, leading to the *in vitro* production of the deacylated **4** (**Fig. 2c**). In contrast, no conversation was detected in the control experiment (*E. coli* carrying empty vector) (**Fig. 2c**). To further support this observation, we inserted a copy of *mucF* into the secondary expression site of pEXT06, resulting in the plasmid pEXT07. Its expression in *E. coli* BAP1 further led to the formation of **4** (**Fig. 2b**). Collectively, these mutations, *in vitro*, and *in vivo* expression studies support the hypothesis that MucF is a newly recognized deacylase responsible for converting **1**-**3** to **4**. Notably, MucF showed sequence similarity to a large group of hypothetical proteins from the genomes of bacteria associated with the human gut and skin (**Supplementary Fig. 33**). We therefore speculate MucF joins a large family of unrecognized deacylases that may play important roles within the human microbiota.

Next, we took two approaches to determine whether RTCs and MUC play roles in mediating inter-species bacterial competition. Frist, we used a plate-based competition assay, in which a colony of *S. mutans* UA159 (a model organism for caries disease), *S. mutans* B04Sm5 (producing **1**-**4**), *S. mutans* B04Sm5/*ΔmucD* (**1**-**4** deficient strain), or *S. mutans* B04Sm5/*ΔmucF* (producing **1**-**3** exclusively) was plated next to nascent colonies of other oral bacteria, including *Rothia mucilaginosa, S. sanguinis, S. gordonii, S. mitis, S. pneumoniae*, and *S. salivarius* (**Fig. 3**). In general, *S. mutans* B04Sm5 exhibited greater inhibition of neighbors than the *S. mutans* model strain UA159. In contrast, *S. mutans* B04Sm5/*ΔmucD*, which lacks the production of compounds **1**-**4**, dramatically exhibited reduced inhibition of its neighbors. For instance, *S. mutans* B04Sm5 impaired the growth of *S. sanguinis* completely, while *S. mutans* B04Sm5/*ΔmucD* only showed slight inhibition of *S. sanguinis*. Furthermore, although the growth of *S. mutans* B04Sm5/*Δmuc*F was impaired by overproducing **1**-**3**, it exhibited the strongest inhibition of neighbors among all tested strains. *S. sanguinis* is one of the predominant species of the indigenous oral biota colonizing dental plaque, which is normally associated with healthy dental biofilm(32). The antagonistic relationship between *S. sanguinis* and *S. mutans* is well-characterized, and plays an important role in caries development(33, 34). Therefore, we determined the Minimum Inhibitory Concentration (MIC) of isolated **1** and **4** against the *S. sanguinis* ATCC 49296. Remarkably, we observed significant antibacterial activity of **1** against the *S. sanguinis* (MIC=3.1 µM), However, **4** did not show any antibacterial activities against *S. sanguinis* up to a concentration of 2 mM. These results provide compelling support that the *muc* pathway, through compounds **1-3**, provides a competitive advantage for *S. mutans* B04Sm5 by inhibiting the growth of its competitors. The increased competitive fitness conferred by *muc* is likely to increase the virulence *S. mutans* strains bearing the gene cluster. As *S. mutans* is an exceptionally productive biofilm-former, higher numbers of *S. mutans* are likely to increase plaque biofilm formation and promote the dysbiosis which leads to the formation of caries lesions. Interestingly, Chen and colleagues showed that **4** can significantly suppress the infiltration of leukocytes (CD45^+^ cells) into the Matrigel plug in a mice model, suggesting an anti-inflammatory activity(23).

**Figure 3.**
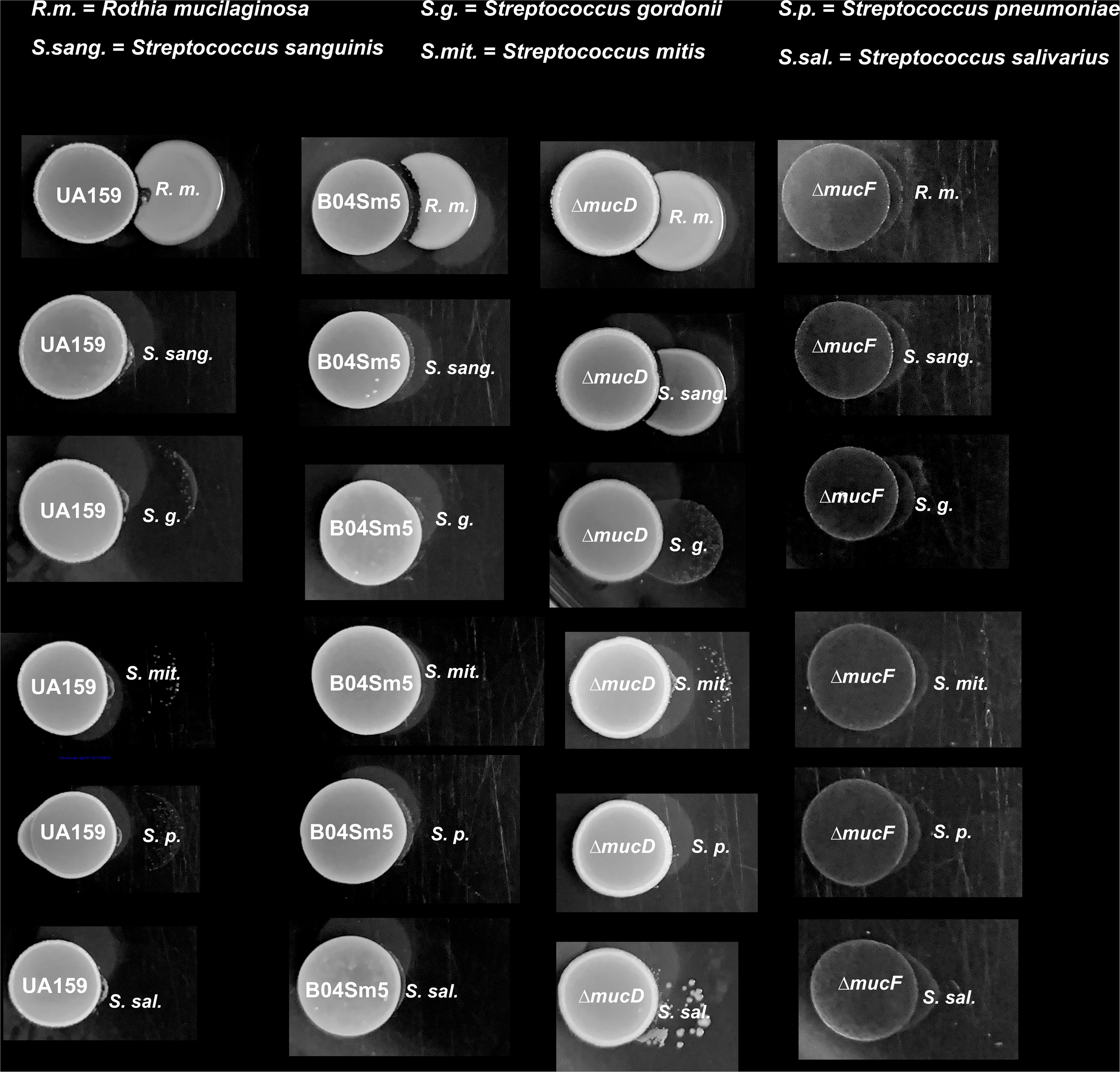
Interspecies competition assay. Overnight cultures of oral commensal bacteria: *R. mucilaginosa, S. gordonii*, and *S. sanguinis, S. mitis, S. pneumoniae*, and *S. salivarius* were plated next to the indicated *S. mutans* strain (UA159, B04Sm5, Δ*mucD*, Δ*mucF*)).

In summary, we describe a versatile biosynthetic pathway from an oral pathogen *S. mutans* B04Sm5, which can produce three types of compounds with divergent biological activities. These include three *N*-acyl tetramic acids (**1**-**3**) that display antibacterial properties against oral commensal bacteria, a new organic acid (**4**) with a reported anti-inflammatory activity in a mice model(23), and a previously characterized SDSF with the ability to interact with pathogenic oral fungi(31). Although two antibiotics have been discovered from the commensal bacteria of human, including lactocillin(13) and lugdunin(35), RTCs, to our knowledge, are the first group of low molecular weight antibacterial molecules identified from a human opportunistic pathogen. While this study merely scrapes the “tip of the iceberg” of the recently identified oral biosynthetic potential(13, 36), these findings clearly exemplify that deeper exploration of leads provided by genome mining studies will help elucidate the complex ecological underpinnings of the human microbiome and its relationship to disease.

## METHODS

### General methods

A complete list of the primers, plasmids, and strains used in this study can be found in **Supplementary Table 6**. PCR products were amplified with PrimeSTAR HS DNA polymerase (Clontech Laboratories, Inc., USA). DNA isolations and manipulations were carried out using standard protocols. *Escherichia coli* strains were cultivated in LB medium (Thermo Fisher Scientific, USA) supplemented with appropriate antibiotics. *S. mutans* B04Sm5 and its respective derivatives were all grown on Brain Heart Infusion (BHI) agar or liquid medium (BD Biosciences, USA) at 37 °C in a CO_2_ incubator (5% CO_2_/95% air). *Lactobacillus reuteri* LTH2584 was also grown on MRS medium (BD Biosciences, USA) or agar at 37 °C in a CO_2_ incubator (5% CO_2_).

### Production, extraction, and detection of reutericyclins (1-3) and mutanocyclin (4) from *S. mutans*

50 milliliters of BHI medium were inoculated with a loop of glycerol stock of *S. mutans* or a mutant thereof overnight. Thirty milliliters of preculture was inoculated into 3 liters of BHI medium containing 1% glucose. After 12 hours incubation, 60 g of autoclaved Amberlite XAD7-HP resin (Sigma-Aldrich, USA) was added to the cultures. The cultures were incubated for another 36 hours at 37 °C in a CO_2_ incubator (5% CO_2_). The resin was recovered and washed twice with water (200 mL) and the extract was subsequently extracted twice with ethyl acetate (400 mL in total). The organic phase was evaporated, and extracts were dissolved in 2 mL methanol. Each extract was monitored at 280 nm during separation by HPLC using a Kinetex® C18 100 Å, LC Column (5 μm, 150 × 2.1 mm; Phenomenex, US) as follows: 0-15 min, 30% B; 15-16 min, 30%-100% B; 16-25 min, 100% B; 26-27 min, 100%-30% B; 28-35 min, 30% B (solvent A: H_2_O/TFA (999:1, v/v); solvent B: CH_3_CN/TFA (999:1)).

### Construction of *S. mutans* knockout plasmids

A 1010-bp fragment containing the spectinomycin resistance gene (*spec*^R^) was amplified from pCAPB2(37) with primers spec_fwd and spec_rev (**Supplementary Table 6**). The left (532 bp) and right (555 bp) flanks of *mucD* were amplified from the genomic DNA of *S. mutans* B04Sm5 with the primer pairs of mucD_KO_L-fwd/mucD_ KO_L-rev and mucD_ KO_R-fwd/mucD_ KO_R-rev (**Supplementary Table 6**), respectively. These three PCR products were assembled with a double digested pUC19 (PstI and EcoRI) using a NEBuilder HiFi DNA Assembly kit (New England Biolabs, USA), which resulted in the vector pEXT01. Amplification of the left (602 bp) and right (611 bp) homology arms for knocking out *mucF* were accomplished with primer pairs mucF_ KO_L-fwd/mucF_ KO_L-rev and mucF_ KO_R-fwd/mucF_ KO_R-rev (**Supplementary Table 6**), respectively. These two PCR products and *spec*^R^ cassette were further cloned into pUC19 to give pEXT02 using the method described above. Vector clones were verified by restriction analysis and sequencing.

### Generation of Δ*mucD* and Δ*mucF* mutants

The disruption cassettes were amplified from pEXT01 (2159 bp) and pEXT02 (2159 bp) using primer pairs mucD_KO_L-fwd/mucD_ KO_R-rev and mucF_ KO_L-fwd/mucF_ KO_R-rev (**Supplementary Table 6**), respectively. PCR products were digested by DpnI and then purified using the QIAquick PCR Purification Kit (Qiagen, USA). The disruption cassettes were transferred to *S. mutans* B04Sm5 by a previously reported protocol(38). Spectinomycin resistance clones were selected by growth on BHI agar supplemented with 500 μg/mL spectinomycin, confirmed by PCR and sequencing, and designated as *S. mutans* B04Sm5/Δ*mucD* and *S. mutans* B04Sm5/Δ*mucF*.

### Generation of *muc* expression plasmid

The 8.6-kb DNA region containing *mucA-E* was PCR amplified from the genomic DNA of *S. mutans* B04Sm5 in two fragments (each approximately 4 kb) with primer pairs mucA-E_ fwd1/ mucA-E_rev1 and mucA-E_fwd2/ mucA-E_rev2 (**Supplementary Table 6**). These fragments were cloned into the XhoI site of pACYCDuet-1 by a NEBuilder HiFi DNA Assembly kit (New England Biolabs, USA), resulting in the plasmid pEXT06. To construct pEXT07, *mucF* was amplified with primers mucF-coexp_fwd and mucF_coexp_rev. PCR product and pEXT06 were digested with the restriction enzyme pair NcoI/BamHI and ligated with T4 DNA ligase (New England Biolabs, USA). The resulting vectors were verified by restriction analysis and sequencing. pEXT06 and pEXT07 were transformed into *E. coli* BAP1, respectively.

### Expression, extraction, and detection of *muc* expression in *E. coli* BAP1

*E. coli* BAP1(39) containing pEXT06 or pEXT07 were cultivated on LB plates supplemented with 1% glucose and 50 μg/mL chloramphenicol at 37 °C. The following day, a loop of *E. coli* cells was transferred for precultures grown at 37 °C in 10 mL LB medium supplemented with 1% glucose and 50 μg/mL chloramphenicol for 4-5 h. One microliter of each preculture was transferred to 50 mL of fresh LB with the same supplements and grown at 37 °C to an OD_600_ of 0.4 to 0.6. Cultures were induced with 200 µM IPTG and incubated for an additional 12-14 h at 30°C with shaking (220 rpm). Cultures were harvested and 1 mL of H_2_O supplemented with 0.5 mg/mL lysozyme was added to the pellets. Cells were disrupted by sonication at room temperature. The lysates were acidified with acetic acid (1% final concertation) and extracted by twice with an equal volume of EtOAc. The organic phase was evaporated, resuspended in MeOH (0.2 mL), and filtered through Acrodisc MS PTFE Syringe filters (Pall Inc., Ann Arbor, MI, USA) prior to HPLC analysis. Each extract was monitored at 280 nm during separation by HPLC using a Kinetex® C18 100 Å, LC Column (5 μm, 150 × 2.1 mm; Phenomenex, US) as follows: 0-10 min, 30% B; 10-11 min, 30%-100% B; 11-25 min, 100% B; 26-27 min, 100%-30% B; 28-35 min, 30% B (solvent A: H_2_O/TFA (999:1, v/v); solvent B: CH_3_CN/TFA (999:1)).

### Feeding experiments for biosynthetic pathway study

*S. mutans* B04Sm5 was cultivated in 50 mL BHI medium supplemented with 1% glucose and 200 mg/L isotope-labeled [^13^C_1_] *L*- or *D*-leucine (Sigma-Aldrich, USA). The compounds were extracted and isolated using the method described above. Each extract prepared was dissolved in 100 µL MeOH for MS analysis using the method described in the supplementary information (**Supplementary Note 1**).

### Expression and activity mensuration of MucF in *E. coli*

Primer pair mucF_pET_fwd/mucF_ pET_rev (**Supplementary Table 6**) was used for amplification of *mucF* from the genomic DNA of *S. mutans* B04Sm5. The PCR product was cloned into the NcoI and XhoI sites of pET28a to obtain pEXT26 (with a C-terminal His-tag). Next, pET28a and pEXT26 were transferred into *E. coli* Rosetta2™(DE3)pLys, respectively. Single clones were picked for precultures grown overnight at 37°C in TB broth (Thermo Fisher Scientific, USA) with 50 μg/mL kanamycin and 50μg/mL chloramphenicol at 37°C. One microliter of preculture was transferred to 1 L of fresh TB broth with the same antibiotics and grown at 37°C to an OD600 of 0.4 to 0.6. Cultures were induced with 500 µM IPTG and incubated for an additional 16 h at 18°C with shaking (220 rpm). Cultures were harvested and 10 mL of buffer (50 mM Tris-HCl, pH 8, 150 mM NaCl, 10% glycerol) supplemented with 0.5 mg/mL lysozyme and 0.5 mM PMSF was added to the pellets. Cells were disrupted by sonication at 4°C. The lysate was used for MucF activity testing. The assay mixture for the reaction (100 μL) consisted of 96 μL *E. coli* lysate (both carrying empty pET28a or pEXT26) and 4 μl reutericyclin A (**1**) solution (6.6 mM, 80% EtOH). The reaction solutions were prepared on ice and incubated at 37°C for 10 min, 30 min and 60 min. Reactions were terminated by the addition of 1 μL acetic acid and then extracted twice with 200 μL EtOAc. After centrifugation of the assay at 12,000 *g* for 10 min, the organic phase was evaporated and resuspended in 100 μL MeOH (0.2 mL). The extracts were monitored by HPLC.

### Agar plate-based assays

The interspecies competition assays with *S. mutans* (UA159, B04Sm5, Δ*mucD*, Δ*mucF*), *R. mucilaginosa, S. gordonii*, and *S. sanguinis, S. mitis, S. pneumoniae*, and *S. salivarius* were performed as described previously(40), with some modifications. 8 µl of overnight cultures of *S. mutans* strains were inoculated onto BHI agar and incubated at 37°C under a 5% CO_2_/95% air atmosphere (vol/vol). After a 24 h incubation period, an overnight culture of the indicated competing species was inoculated next to the *S. mutans* and the plates were incubated for an additional 24 h and subsequently photographed.

## Acknowledgments

The authors thank Y. Li (NYU College of Dentistry, USA) for providing *S. mutans* strain B04Sm5, M.G. Gänzle (University of Alberta, Canada) for providing *L. reuteri* LTH2584 strain, C. Khosla (Stanford University, USA) for *E. coli* BAP1, and J.J. Zhang (Massachusetts Institute of Technology, USA), M.S. Donia (Princeton University, USA) and M.A. Fischbach (Stanford University, USA) for valuable discussions. This work was supported by NIH grants R00-DE0245543 and R21-DE028609-01 to A.E., R01-GM085770 to B.S.M., and F32-DE026947 to J.B., and the Japan Society for Promotion of Science Overseas Research Fellowship to Y.K.

## Author contributions

X.T. and A.E. designed the research and X.T. analyzed the *muc* pathway. X.T. generated and analyzed the mutants, performed the biochemical experiments and the heterologous expression experiments. Y. K. and X.T. purified the compounds and elucidated the structures of all compounds. X.T., P.A.J. and S.M.K.M. performed the chemical synthesis. X.T., Y.K., J.G. and T.H. performed mass spectrometry experiments and analyzed mass spectrometry data. X.T., J.B. and S.L. designed and performed the agar plate-based assays. X.T., Y.K., J.B., A.E. and B.S.M. wrote the manuscript. All authors analyzed and discussed the data and contributed to the writing of the manuscript.

