## Supplementary information for "Cariogenic *Streptococcus mutans* produces strain-specific antibiotics that impair commensal colonization"

### Supplementary Note 1

#### Purification and structure elucidation of reutericyclins A-C (1-3).

The crude extract was prepared as described above from 3 liters of culture, and then purified using a Luna C18 (2) column (5 μm, 100 × 21.2 mm, Phenomenex, USA) with following conditions; $CH_3CN/H_2O$/TFA (85:15:0.1, v/v/v), 10 mL/min, UV detector at λ = 210 and 280 nm, along with the Agilent Technologies system composed of a PrepStar pump, a ProStar 410 autosampler, and a ProStar 325 UV detector (Agilent Technologies, Inc., USA). Collected fractions containing reutericyclins were evaporated *in vacuo*. After acetonitrile was evaporated, the resultant cloudy aqueous solution was extracted with 1 volume of ethyl acetate twice, and then organic layer was evaporated to obtain pure reutericyclin A (**1**, 5.0 mg), reutericyclin B (**2**, <0.1 mg), and reutericyclin C (**3**, 1.8 mg).

The molecular formula of reutericyclin A (**1**, $C_{20}H_{31}NO_4$) determined from the HR-ESI-MS peak a *m/z* 350.2337 ($[M+H]^+$, calcd. 350.2326, err. 3.1 ppm, **Supplementary Fig. 2**). The structure was confirmed by NMR measurements; $^1$H-NMR, $^{13}$C-NMR, gradient COSY, gradient HSQC and gradient HMBC ($^{2,3}J_{CH}$ = 8 Hz) in $CDCl_3$. Reutericyclin A exists as tautomers of ***a*** and ***b***/***c***, as described in the first isolation paper (1), and ***a***:***b***/***c*** ratio in $CDCl_3$ was estimated to be 10 : 7 by $^1$H NMR spectroscopy. All the proton and carbon signals were assigned based on the 2D NMR analysis (**Supplementary Table 2** and **Figs. 3-7**). The structure of **1** was further comfired by $^1$H NMR and $^{13}$C-NMR comparison with a synthetic standard (**Supplementary Fig. 8**).The absolute configuration of **1** was determined by HPLC using a chiral LC column (Lux® 5 μm i-Amylose-1, 100 x 4.6 mm, Phenomenex, US) in comparison with the reported reutericyclin from *Lactobacillus reuteri* LTH2584 (1) and a synthetic standard (**Supplementary Fig. 9**). The elution condition was $CH_3CN/H_2O$ (85:15, each containing 0.1% TFA) at a flow rate of 1.0 mL/min.

The formula ($C_{20}H_{33}NO_4$) of reutericyclin B (**2**) was determined through HR-ESI-MS measurement ($[M+H]^+$ calcd. 352.2482, found. 352.2498, err. 4.5 ppm). MS/MS analysis suggested the presence of the tetramic acid core (*m/z* 198.1132) and a decanoyl substituent (C10) at the N-1 position (**Supplementary Fig. 1**). Due to the material limitation (less than 0.1 mg), NMR measurements were not performed for **2**.

Structure elucidation of reutericyclin C (**3**), (*E*)-dodec-2-enoyl-reutericyclin, was primarily achieved via HR-ESI-MS and NMR measurements. A molecular formula of **3**, $C_{22}H_{35}NO_4$ ($[M+H]^+$ calcd. 378.2639, found. 378.2661, err. 5.8 ppm), and fragment ion at *m/z* 198.1125 ($C_{10}H_{16}NO_3$) in the MS/MS spectrum indicated the presence of a dodecenoyl substituent (C12) at the N-1 position in **3 (Supplementary Fig. 2)**. All the proton and carbon signals were assigned based on the 2D NMR analysis in $CDCl_3$ (**Supplementary Table 3 and Supplementary Figs. 10-14**). NMR analysis clarified *N*-(*E*)-dodec-2-enoyl structure and the core mutanic acid structure in **3**. Compound **3** was also observed as a mixture of tautomers of ***a*** and ***b***/***c*** in $CDCl_3$.

NMR spectra were recorded on a 600 MHz Bruker Avance III NMR spectrometer (Topspin 2.1.6 software, Bruker) with a 5.0 mm cryoprobe at 20 °C. The solvent signal at 7.26 ppm in the $^{1}H$ NMR spectra and $^{13}CDCl_3$ at 77.16 ppm in the $^{13}C$ NMR spectra were used as internal references. Direct infusion-mass spectrometry (MS) analysis was conducted to collected MS/MS spectra on a Bruker Impact II mass spectrometer (Billerica, MA, U.S.A). Direct injection flow rate was set as 180 µL/h. Compounds were diluted in $CH_3CN/H_2O$ (1:1, v/v). MS conditions for the MS/MS spectra generation were set as follows: capillary voltage, 4500; nebulizer gas flow, 1.6 Bar; dry gas, 6.0 L/min at 220 ºC; funnel 1 RF 150 Vpp; funnel 2 RF, 200 Vpp; isCID energy, 0 eV; hexapole RF: 50 Vpp; Quadrupole ion energy, 4 eV; low mass 50 m/z; collision cell energy, 20 – 50 eV; pre pulse storage 5.0 µs; collision RF, ramp from 350 to 800 Vpp; transfer time ramp from 50 to 100 µs; detection mass range 25 to 1000 m/z; spectra collection rate 2.0 Hz. Each MS/MS spectrum was acquired for 30 seconds.

**Purification and structure elucidation of mutanic acid (4).**

Five liters of *S. mutans* culture was obtained using the same fermentation method mentioned above. The supernatant was adjusted to pH 4.0 and subsequently extracted with an equal volume of EtOAc. The organic phase was evaporated, resuspended in MeOH (5 mL), and filtered through Acrodisc MS PTFE Syringe filters (Pall Inc., Ann Arbor, MI, USA), followed by passage over a reversed-phase C18 silica gel column (50g silica gel, Merck, 70–230 mesh). The polar fraction was removed by elution with 250 mL $CH_3CN/H_2O$/TFA (5:95:0.1, v/v/v). Then, the column was eluted sequentially using 250 mL $CH_3CN/H_2O$/TFA (30:70:0.1, v/v/v) and 250mL $CH_3CN/H_2O$/TFA (70:30:0.1, v/v/v). All fractions were measured by analytical HPLC. The fractions containing mutanic acid were pooled together and evaporated. The sample was resuspended in MeOH (1 mL) prior to semi-preparative HPLC separation on a Phenomenex Luna C18 reversed-phase HPLC column (5 μm, 250 mm × 10 mm) with isocratic elution (30% $CH_3CN$–70% $H_2O$, each solvent contains 0.1% TFA; flow rate 2.5 mL/min), yielding 1.5 mg of mutanic acid (**4**).

The formula ($C_{10}H_{15}NO_3$) of **4** was determined through HR-ESI-MS measurement ($[M+H]^+$ calcd. 198.1125, found. 198.1124, 0.5 ppm). High-Resolution LCMS/MS analysis also revealed the structure **4** is identical to the tetramic acid core in **1-3** (**Supplementary Fig. 2**). The structure of **4** was further comfired by $^1H$ NMR and HPLC comparison with a synthetic standard (**Supplementary Figure 15**). The stereochemistry of **4** was determined by HPLC using a chiral LC column (Lux® 5 µm i-Amylose-1, 100 x 4.6 mm) in comparison with synthetic standards (**Supplementary Figure 16**). The elution condition was $CH_3CN/H_2O$ (40:60, each containing 0.1% TFA) at a flow rate of 1.0 mL/min.

### Synthesis of mutanic acid (4) and reutericyclin A (1).

(*R*)-**4** and (*S*)-**4** were synthesized according to a previous report(2), however, due to the limited supply of *N*-(hydroxysuccinamidyl) acetoacetate, we adapted the procedure using 2,2,6-trimethyl-4H-1,3-dioxin-4-one in xylene (**7**)(3).

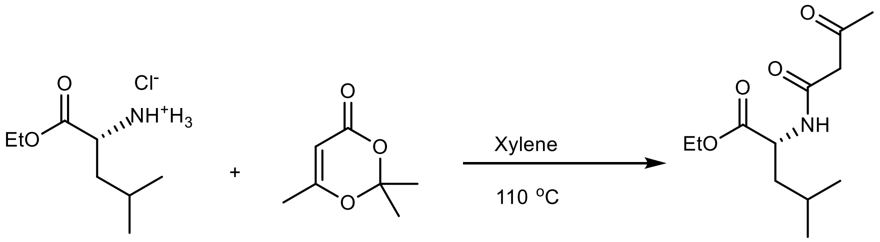

**6** **7** **8**

***Ethyl N-(Acetoacetyl)-D-leucinate*** (**8**): To a solution of D-leucine ethyl ester hydrochloride (**6**, 1.03 mmol, Sigma-Aldrich, US) in xylene (10 mL) at r.t. was added NaOH (408 mg, 1.03 mmol) and stirred for 15 min. Then, 2,2,6-trimethyl-4H-1,3-dioxin-4-one (**7**, 1.7 mL, 1 mmol) was added through a syringe over 15 min. Under $N_2$ gas, the reaction was stirred at 110 °C for another 45 min. The solution was poured into a separatory funnel and washed with 5% HCl solution (4×10mL). The aqueous phase was extracted with $CH_2Cl_2$ (3×40 mL), and the combined organic phases were washed with brine and dried over $MgSO_4$. The solvent was removed, and the residue was purified by flash column chromatography (hexanes/EtOAc 1:2, Alfa Aesar silica gel, 60 Å pore size) to give 1.3 g of ethyl N-(acetoacetyl)-D-leucinate as a pale-yellow oil (53% yield) with a $R_f$ of 0.5 (1:2, hexanes/EtOAc). $^1$H-NMR and $^{13}$C-NMR matched those reported in the literature(2).

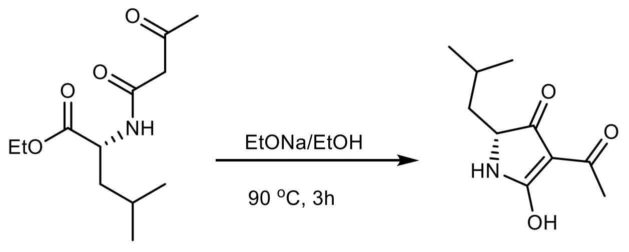

8  (*R*)-**4**

***(2R)-4-Acetyl-1,2-dihydro-5-hydroxy-2-(2-methylpropyl)-3H-pyrrol-3-one*** **((*R*)-4 or (*R*)-mutanic acid)**: (*R*)-**4** was synthesized using a published method(2). 3.9 mL EtOH (200 proof, molecular biology grade), followed by Na metal (0.127 g, 1.07 equivalents) were added to a flame-dried round bottom flask fitted with a dry nitrogen gas line, and were allowed to dissolve over the course of several minutes. In a separate flask, ethyl N-(acetoacetyl)-D-leucinate (1.3 g, 23.5 mmol) was dissolved in 6.8 mL EtOH then transferred dropwise via syringe (over 25 min) to the EtONa solution. The reaction was then heated at reflux for 3h under nitrogen. After cooling to room temperature, the mixture was neutralized with 10% acetic acid in EtOH (1.04 mL), and the solvent was evaporated. The pale-yellow, gel-like residue was dissolved in EtOAc (5 mL) and washed with 1% aqueous acetic acid (3× 5 mL) to remove AcONa. The combined aqueous washes were then extracted with EtOAc (3 × 15 mL), the combined organic phases were dried over $Na_2SO_4$, and the solvent was removed using a rotary evaporator. A portion of the resulting solid residue was recrystallized from hot EtOH to yield 180 mg (17% yield) of (*R*)-mutanic acid ((*R*)-**4**). Further recrystallizations, to yield more material, were not pursued. *Mr* 197.23 g/mol. [*α*]22/D =+103° (c=0.1, EtOH), lit. = +117°). $^1$H-NMR and $^{13}$C-NMR matched those reported in literature(2). See included $^1$H-NMR, $^{13}$C-NMR, COSY, HSQC and HMBC spectra (**Supplementary Figs. 17-21**). HPLC analysis of the synthetic compound using a chiral LC Column (Lux® 5 µm i-Amylose-1, 100 x 4.6 mmon) showed two fully separated peaks (**Supplementary Fig. 22**) corresponding to a R/S mixture of 80.5 : 19.5 (61.2% ee).

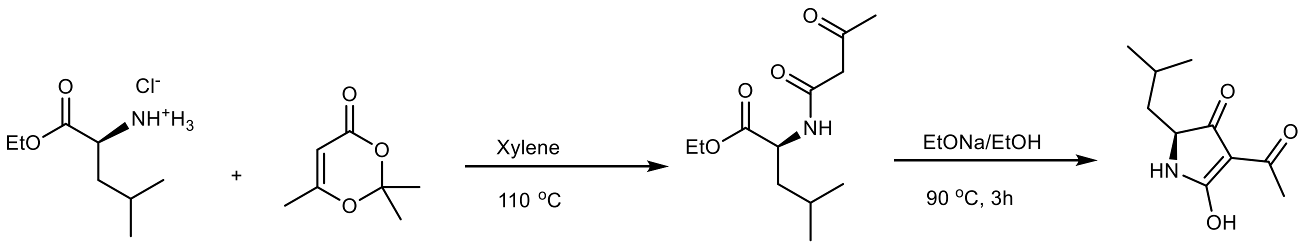

***(2S)-4-Acetyl-1,2-dihydro-5-hydroxy-2-(2-methylpropyl)-3H-pyrrol-3-one*** **((*S*)-4 or (*S*)-mutanic acid**): (*S*)-**4** was synthesized following the same procedure as described in the synthesis of (*R*)-**4**, except L-leucine methyl ester hydrochloride (**9**, Sigma-Aldrich, USA) was used to prepare ethyl N-(acetoacetyl)-L-leucinate (**10**). HPLC analysis of the synthetic compound using a chiral LC Column (Lux® 5 µm i-Amylose-1, 100 x 4.6 mmon) showed two fully separated peaks (**Supplementary Fig. 22**) corresponding to a R/S mixture of 38.6 : 61.4 (22.7% ee). [α]22/D =-25° (c=0.1, EtOH).

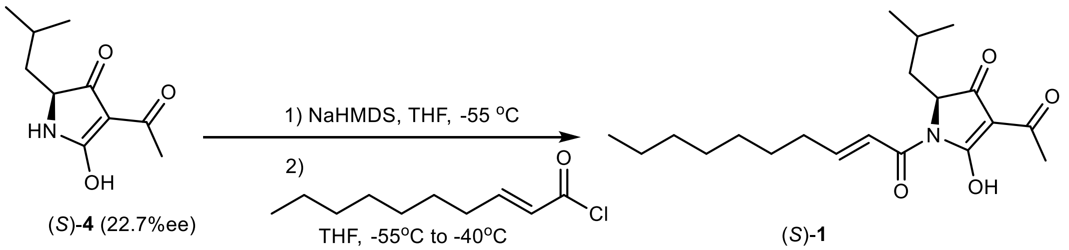

***(2S)-4-Acetyl-1,2-dihydro-5-hydroxy-2-(2-methylpropyl)-1-[(2E)-1-oxodec-2-enyl]-3H-pyrrol-3 one*** (**1**, Reutericyclin A). **1** was synthesized using a published method (4). To an ice-cold solution of (*E*)-dec-2-enoic acid (449 µl, 2.31 mmol) in 4 mL of dry DCM (with 2 drops of DMF) maintained under an argon atmosphere was added oxalyl chloride (208 µL, 2.42 mmol). After stirring at room temperature for 2h, the solvent was evaporated and the (*E*)-dec-2-enoyl chloride was dissolved in 5mL of dry THF. In another flask maintained under an argon atmosphere, a solution of (*S*)-**4** (100 mg, 0.507 mmol, 22.7% ee) in 5mL of dry THF was cooled to -55°C and sodium bis(trimethylsilyl)amide (1M in THF, 223 mg, 1.22 mmol) was added. After stirring for 10 min, 2.2 mL of the (*E*)-dec-2-enoyl chloride in THF was added dropwise over a period of 15 mins. After the addition, the temperature was raised from -55°C to -40°C over 30 mins and then maintained at -40°C for another 30 mins. The reaction was quenched by the addition of acetic acid (87 µL, 1.52 mmol). The solvent was evaporated, and the crude mixture was separated by reverse phase chromatography to provide 30 mg (17%) of product. (1:0.7 mixture of tautomers)

$^{1}$**H-NMR** (500 MHz, $CDCl_3$, both tautomers): δ 7.29 (dt, *J* = 15.8, 1.5 Hz, 1H), 7.24 – 7.09 (m, 2H), 4.71 (dd, *J* = 8.2, 3.1 Hz, 1H), 4.47 (dd, *J* = 7.3, 4.0 Hz, 1H), 2.57 (s, 3H), 2.55 (s, 2H), 2.28 (qd, *J* = 8.2, 7.7, 3.6 Hz, 4H), 1.93 – 1.76 (m, 5H), 1.49 (tt, *J* = 7.5, 3.7 Hz, 4H), 1.36 – 1.23 (m, 16H), 0.96 (d, *J* = 6.0 Hz, 3H), 0.94 (d, *J* = 6.2 Hz, 2H), 0.92 – 0.85 (m, 11H).

$^{13}$**C-NMR** (125 MHz, $CDCl_3$): δ 198.3, 195.0, 193.8, 188.6, 173.7, 166.3, 165.6, 165.0, 152.1, 151.2, 123.1, 123.1, 105.2, 102.7, 63.5, 59.5, 39.5, 39.5, 32.9, 32.9, 31.9, 29.4, 29.3, 29.2, 28.3, 28.2, 24.6, 24.5, 23.7, 23.7, 22.9, 22.8, 22.4, 22.3, 20.3, 14.2.

**Supplementary Table 1**. Sequenced strains with *muc* pathways distributed globally.

| Strain Name | Genbank Accession | Country of Origin | Associated Disease |
|---|---|---|---|
| *S. mutans* B04Sm5 | ALYY00000000 | USA | Childhood caries |
| *S. mutans* B05Sm11 | ALYO00000000 | USA | Childhood caries |
| *S. mutans* B06Sm2 | ALZA00000000 | USA | Childhood caries |
| *S. mutans* B107SM-B | ALYS00000000 | USA | Childhood caries |
| *S. mutans* B082SM-A | ALYZ00000000 | USA | Childhood caries |
| *S. mutans* AC4446 | AOCA00000000 | Germany | Infective endocarditis |
| *S. mutans* KK23 | AOBZ00000000 | Germany | Childhood caries |
| *S. mutans* NN2025 | NC_013928 | Japan | Dental caries |
| *S. mutans* B07Sm2 | ALYT00000000 | USA | no |
| *S. mutans* B115SM-A | ALZH00000000 | USA | no |
| *S. mutans* G123 | AHSC00000000 | UK | N/A |
| *S. mutans* M21 | AHSD00000000 | UK | N/A |
| *S. mutans* NLML4 | AHSH00000000 | UK | N/A |
| *S. mutans* NLML9 | AHSJ00000000 | UK | N/A |
| *S. mutans* M2A | AHSK00000000 | UK | N/A |
| *S. mutans* W6 | AHSO00000000 | UK | N/A |
| *S. mutans* 14D | AHSY00000000 | Iceland | N/A |
| *S. mutans* B | AHTB00000000 | Iceland | N/A |
| *S. mutans* 24 | AHTE00000000 | Iceland | N/A |
| *S. mutans* 11VS1 | AHRT00000000 | Brazil | N/A |
| *S. mutans* SM6 | AHSR00000000 | China | N/A |
| S. *macacae* ATCC35911 | AEUW00000000 | Dental plaque of monkey | |

**Supplementary Table 2.** NMR spectroscopic data of compound **1**, reutericyclin A, in $CDCl_3$ (600 MHz).

| | **1** (tautomer ***a***) | | | **1** (tautomers ***b***/***c***) | |
|---|---|---|---|---|---|
| position | $\delta_{C,}$ type | $\delta_H$ (*J* in Hz) | gHMBC | $\delta_{C,}$ type | $\delta_H$ (*J* in Hz) |
| 2 | 166.3, C | | | 173.7, C | |
| 3 | 105.1, C | | | 102.7, C | |
| 4 | 198.3, C | | | 193.9, C | |
| 5 | 59.5, CH | 4.71, dd (8.2, 3.2) | C2, C3, C4, C6 | 63.5, CH | 4.47, dd (7.3, 4.0) |
| 6 | 39.5, $CH_2$ | 1.91, m | C4, C5 | 39.4, $CH_2$ | 1.80, m |
| | | 1.80, m | C4, C5 | | |
| 7 | 24.6, CH | 1.86, m | C6 | 24.5, CH | 1.86, m |
| 8 | 22.3, $CH_3$ | 0.96, d (6.1) | C6, C7, C9 | 22.4, $CH_3$ | 0.94, d (6.1) |
| 9 | 23.7, $CH_3$ | 0.90, d (6.6) | C6, C7, C8 | 23.7, $CH_3$ | 0.90, m |
| 10 | 195.1, C | | | 188.7, C | |
| 11 | 23.0, $CH_3$ | 2.57, s | C3, C10 | 20.3, $CH_3$ | 2.55, s |
| 12 | 165.7, C | | | 165.0, C | |
| 13 | 123.1, CH | 7.28, br d (15.4) | C12, C14, C15 | 123.0, CH | 7.22, m |
| 14 | 151.3 CH | 7.15, m | C12, C13, C15, C16 | 152.2, CH | 7.20, m |
| 15 | 32.9, $CH_2$ | 2.28, br p (7.2) | C13, C14, C16 | 33.0, $CH_2$ | 2.28, br p (7.2) |
| 16 | 28.3, $CH_2$ | 1.49, m | C14, C15, C17-C18 | 28.2, $CH_2$ | 1.49, m |
| 17 | 29.3, $CH_2$ | 1.32, m | | 29.3, $CH_2$ | 1.32, m |
| 18 | 29.3, $CH_2$ | 1.30, m | overlapped | 29.3, $CH_2$ | 1.30, m |
| 19 | 31.9, $CH_2$ | 1.26, m | | 31.9, $CH_2$ | 1.26, m |
| 20 | 22.8, $CH_2$ | 1.28, m | | 22.8, $CH_2$ | 1.28, m |
| 21 | 14.2, $CH_3$ | 0.87, t (7.0) | C19, C20 | 14.2, $CH_3$ | 0.87, t (7.0) |

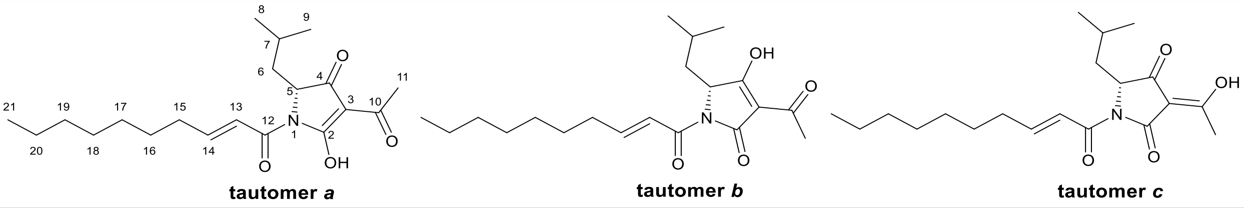

**Supplementary Table 3.** NMR spectroscopic data of compound **3**, reutericyclin C ((*E*)-dodec-2-enoyl-reutericyclin), in $CDCl_3$ (600 MHz).

| | **3** (tautomer ***a***) | | | **3** (tautomers ***b***/***c***) | |
|---|---|---|---|---|---|
| position | $\delta_{C,}$ type | $\delta_H$ ($J$ in Hz) | gHMBC | $\delta_{C,}$ type | $\delta_H$ ($J$ in Hz) |
| 2 | 166.3, C | | | 173.7, C | |
| 3 | 105.2, C | | | 102.7, C | |
| 4 | 198.3, C | | | 193.8, C | |
| 5 | 59.5, CH | 4.71, dd (8.2, 3.0) | C2, C3, C4, C6 | 63.5, CH | 4.47, dd (7.5, 4.0) |
| 6 | 39.5, $CH_2$ | 1.89, m | C4, C5 | 39.5, $CH_2$ | 1.80, m |
| | | 1.80, m | C4, C5 | | |
| 7 | 24.6, CH | 1.86, m | C6 | 24.5, CH | 1.86, m |
| 8 | 22.3, $CH_3$ | 0.96, d (6.5) | C6, C7, C9 | 22.4, $CH_3$ | 0.94, d (6.4) |
| 9 | 23.7, $CH_3$ | 0.91, d (6.2) | C6, C7, C8 | 23.7, $CH_3$ | 0.90, m |
| 10 | 195.0, C | | | 188.7, C | |
| 11 | 23.0, $CH_3$ | 2.57, s | C3, C10 | 20.3, $CH_3$ | 2.55, s |
| 12 | 165.6, C | | | 165.0, C | |
| 13 | 123.1, CH | 7.29, dt (15.4, 1.5) | C12, C14, C15 | 123.1, CH | 7.22, m |
| 14 | 151.2 CH | 7.15, m | C12, C13, C15, C16 | 152.1, CH | 7.20, m |
| 15 | 32.9, $CH_2$ | 2.28, br quin (7.2) | C13, C14, C16 | 33.0, $CH_2$ | 2.28, br quin (7.2) |
| 16 | 28.3, $CH_2$ | 1.49, m | C14, C15, C17-C18 | 28.2, $CH_2$ | 1.49, m |
| 17 | 29.7, $CH_2$ | 1.32, m | | 29.3, $CH_2$ | 1.32, m |
| 18 | 29.6, $CH_2$ | 1.30, m | | 29.3, $CH_2$ | 1.30, m |
| 19 | 29.4, $CH_2$ | 1.30, m | overlapped | 32.0, $CH_2$ | 1.26, m |
| 20 | 29.3, $CH_2$ | 1.30, m | | 32.0, $CH_2$ | 1.26, m |
| 21 | 32.0, $CH_2$ | 1.26, m | | 32.0, $CH_2$ | 1.26, m |
| 22 | 22.8, $CH_2$ | 1.28, m | | 22.8, $CH_2$ | 1.28, m |
| 23 | 14.3, $CH_3$ | 0.88, t (7.2) | C21, C22 | 14.3, $CH_3$ | 0.88, t (7.2) |

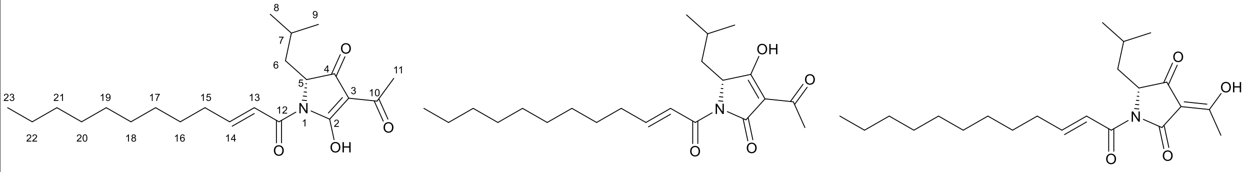

**tautomer *a*** **tautomer *b*** **tautomer *c***

**Supplementary Table 4.** NMR spectroscopic data of compound **5**, reutericyclin D, in $CDCl_3$ (600 MHz).

| | reutericyclin D - a | | | reutericyclin D - b/c | |
|---|---|---|---|---|---|
| position | $\delta_{C,}$ type | $\delta_H$ (J in Hz) | gHMBC | $\delta_{C,}$ type | $\delta_H$ (J in Hz) |
| 1 | N | | | N | |
| 2 | 166.1, C | | | 177.3, C | |
| 3 | 105.0, C | | | 102.4, C | |
| 4 | 198.2, C | | | 193.6, C | |
| 5 | 59.2, CH | 4.64, dd | C2, C4, C6 | 63.2, CH | 4.40, dd |
| 6 | 39.2, $CH_2$ | 1.87, m | C5 | 39.2, $CH_2$ | 1.78, m |
| | | 1.77, m | C4, C5 | | |
| 7 | 24.3, CH | 1.85, m | C6 | 24.3, CH | 1.85, m |
| 8 | 22.2, $CH_3$ | 0.94, d | C6, C7, C9 | 22.2, $CH_3$ | 0.94, d |
| 9 | 23.4, $CH_3$ | 0.90, d | C6, C7, C8 | 23.4, $CH_3$ | 0.90, d |
| 10 | 194.9, C | | | 188.5, C | |
| 11 | 22.7, $CH_3$ | 2.56, s | C3, C10 | 20.0, $CH_3$ | 2.54, s |
| 12 | 173.3, C | | | 173.3, C | |
| 13 | 38.0, $CH_2$ | 2.96, dd | | 38.0, $CH_{11}$ | 2.96, dd |
| 14 | 24.2, $CH_2$ | 1.67, dd | C12, C13, C15 | 24.2, $CH_{12}$ | 1.67, dd |
| 15 | 29.4, $CH_2$ | 1.35, m | | 29.4, $CH_2$ | 1.35, m |
| 16 | 29.4, $CH_2$ | 1.25, m | | 29.4, $CH_2$ | 1.25, m |
| 17 | 29.4, $CH_2$ | 1.25, m | | 29.4, $CH_2$ | 1.25, m |
| 18 | 29.4, $CH_2$ | 1.25, m | overlapped | 29.4, $CH_2$ | 1.25, m |
| 19 | 29.4, $CH_2$ | 1.25, m | | 29.4, $CH_2$ | 1.25, m |
| 20 | 29.4, $CH_2$ | 1.25, m | | 29.4, $CH_2$ | 1.25, m |
| 21 | 31.9, $CH_2$ | 1.25, m | | 31.9, $CH_2$ | 1.25, m |
| 22 | 22.8, $CH_2$ | 1.28, m | | 22.8, $CH_2$ | 1.28, m |
| 23 | 14.1, $CH_3$ | 0.87, t | C21, C22 | 14.1, $CH_3$ | 0.87, t |

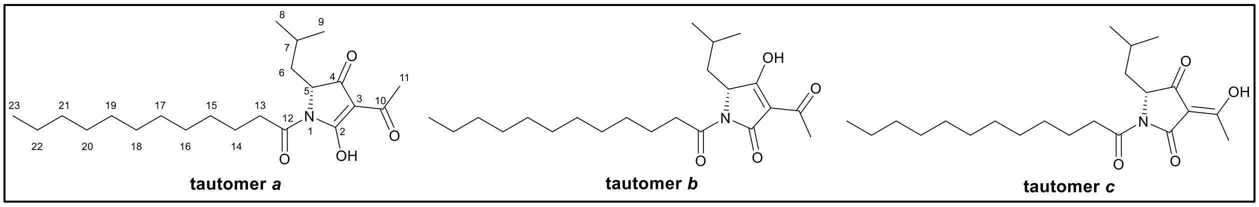

**Supplementary Table 5.** Deduced function of genes from the *muc* gene cluster in *Streptococcus mutans* B04Sm5.

| Gene | AAs | Protein homolog (BLAST) | Identity/ Similarity | Proposed function | Identity to Rtc |
|---|---|---|---|---|---|
| ***mucA*** | 359 | PhlA (AAY95144) | 31%/54% | hydroxymethylglutaryl-CoA synthase | Rtc A, 57% |
| ***mucB*** | 403 | PhlC (5M3K_C) | 36%/54% | thiolase | Rtc C, 66% |
| ***mucC*** | 148 | PhlB (AAM27407) | 38%/58% | hypothetical protein | Rtc B, 69% |
| ***mucD*** | 1024 | Rtc N (WP_035152898) | 48%/65% | NRPS, C-A-T* | Rtc N, 48% |
| ***mucE*** | 913 | Rtc K (AJO68346) | 53%/71% | Type I PKS, KS-T-TE* | Rtc K, 53% |
| ***mucF*** | 167 | WP_099390897 | 57%/76% | HXXEE domain-containing protein | |
| ***mucG*** | 180 | WP_099390896 | 62%/81% | TetR/AcrR family transcriptional regulator | |
| ***mucH*** | 180 | WP_002888975 | 63%/82% | TetR/AcrR family transcriptional regulator | |
| ***mucI*** | 598 | WP_073950866 | 73%/87% | DHA2 family efflux MFS transporter permease subunit | |

*C, condensation; A, adenylation; T, thiolation; KS, ketosynthase; TE, thioesterase.

**Supplementary Table 6**. Plasmids, primers, and strains used in this study

| Strain/Plasmid | Description | Ref |
|---|---|---|
| **Plasmids** | | |
| pUC19 | *amp*$^{R}$, *lacZα*, *lac* promoter, *pMB1 ori* | NEB |
| pCapB2 | *amp*$^{R}$, *spec*$^{R}$, *ARSH4/CEN6*, *pBR322 ori* | (5) |
| pEXT01 | pUC19 derivative carrying the cassette for knocking out *mucD* | This study |
| pEXT02 | pUC19 derivative carrying the cassette for knocking out *mucF* | This study |
| pACYCDuet-1 | The vector encodes two multiple cloning sites (MCS) each of which is receded by a T7 promoter, lac operator and ribosome binding site (rbs) | Novagen |
| pEXT06 | pACYCDuet-1 derivative that carries a 8.6 kb genomic region containing *mucA-D* in the first MCS. | This study |
| pEXT07 | pEXT06 derivative in which *ttmF* has been inserted into the second MCS. | This study |
| pET28a | Empty protein expression vector | Novagen |
| pEXT26 | pET28a derivative carring *ctcF* | This study |
| **Primers** | | |
| spec_fwd | CAATTTTTTTATAATTTTTTTAATCTGTTA | This study |
| spec_rev | ATAACTAATAACGTAACGTG | This study |
| mucD_KO_L-fwd | ACCATGATTACGCCAAGCTTGCATGCCTGCAGATATGGTTTATCAACTGTG | This study |
| mucD_ KO_L-rev | AAGGGCGACACGGAAATGTTGAATACTCATTCTTATTTATCCTTCTTTC | This study |
| mucD_ KO_R-fwd | ATAGGTGCCTCACTGATTAAGCATTGGTAAATATGCAATTATTGGAATGG | This study |
| mucD_ KO_R-rev | AGTCACGACGTTGTAAAACGACGGCCAGTGAATTCAAATCACAAGTATCTGAATC | This study |
| mucF_ KO_L-fwd | ACCATGATTACGCCAAGCTTGCATGCCTGCAGAAAACGACAGATTCTAATGTG | This study |
| mucF_ KO_L-rev | AAATAACAGATTAAAAAAATTATAAAAAAATTGAAATGTAGAAACCATAACCAG | This study |
| mucF_ KO_R-fwd | ATCTCTTGCCAGTCACGTTACGTTATTAGTTATTTTTAACCCTCTTTCATTATTAT | This study |
| mucF_ KO_R-rev | AGTCACGACGTTGTAAAACGACGGCCAGTGAATTCTCTAGCTGAAAATCATTTTC | This study |
| mucA-E_ fwd1 | ATTAGTTAAGTATAAGAAGGAGATATACATATGGGCAATAAACAGATCGG | This study |
| mucA-E_rev1 | CTAATATGACAGTTTTCCGCTG | This study |
| mucA-E_fwd2 | AAGAATATGCCTGAACAGCG | This study |
| mucA-E_rev2 | TTCGCAGCAGCGGTTTCTTTACCAGACTCGAGCCCTAAAATGATATTATTTCCTCTAAG | This study |
| mucF-coexp_fwd | ATTTTGTTTAACTTTAATAAGGAGATATACCATGGATCTATTATCTAATTATTTC | This study |
| mucF_coexp_rev | CGACCTGCAGGCGCGCCGAGCTCGAATTCGGATCCTTAGCCTTTAAATAAGTTTAAC | This study |
| mucF_pET_fwd | GTTTAACTTTAAGAAGGAGATATACCATGGATCTATTATCTAATTATTTCATG | This study |
| mucF_ pET_rev | GGATCTCAGTGGTGGTGGTGGTGGTGCTCGAGGCCTTTAAATAAGTTTAACAG | This study |
| **Strains** | | |
| *E. coli* DH5α | This strain was used to Maintain and propagate vectors | NEB |
| *E. coli* BAP1 | This strain contains the Sfp phosphopantetheinyl transferase and was used for the pathway expression | (6) |
| *E. coli* Rosetta2$^{TM}$ (DE3)pLys | This strains was used for protein expression | Novagen |
| *S. mutans* B04Sm5 | This strain was isolated from a child with severe early childhood caries at NYU College of Dentistry | (7) |
| *S. mutans* B04Sm5/Δ*mucD* | *S. mutans* B04Sm5 derivative, in which *mucD* was replaced by the spectinomycin resistance gene from pCAPB2 | This study |
| *S. mutans* B04Sm5/Δ*mucF* | *S. mutans* B04Sm5 derivative, in which *mucF* was replaced by the spectinomycin resistance gene from pCAPB2 | This study |
| *L. reuteri* LTH2584 | This is the first reported reutericyclin producer isolated from sourdough samples in Germany | (1) |

**Supplementary Figure 1.** Comparison of *muc* and *rtc* gene clusters.

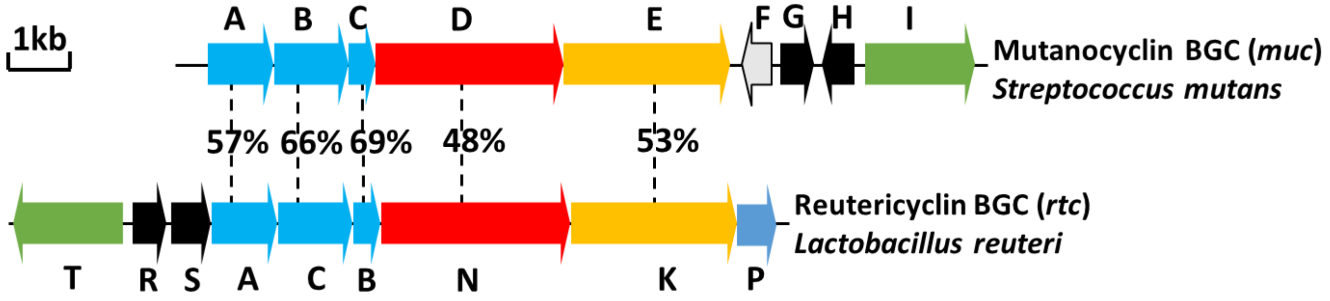

**Supplementary Figure 2**. High-resolution MS/MS spectrometry analysis of compounds **1**-**4** isolated from *S. mutans* B04Sm5.

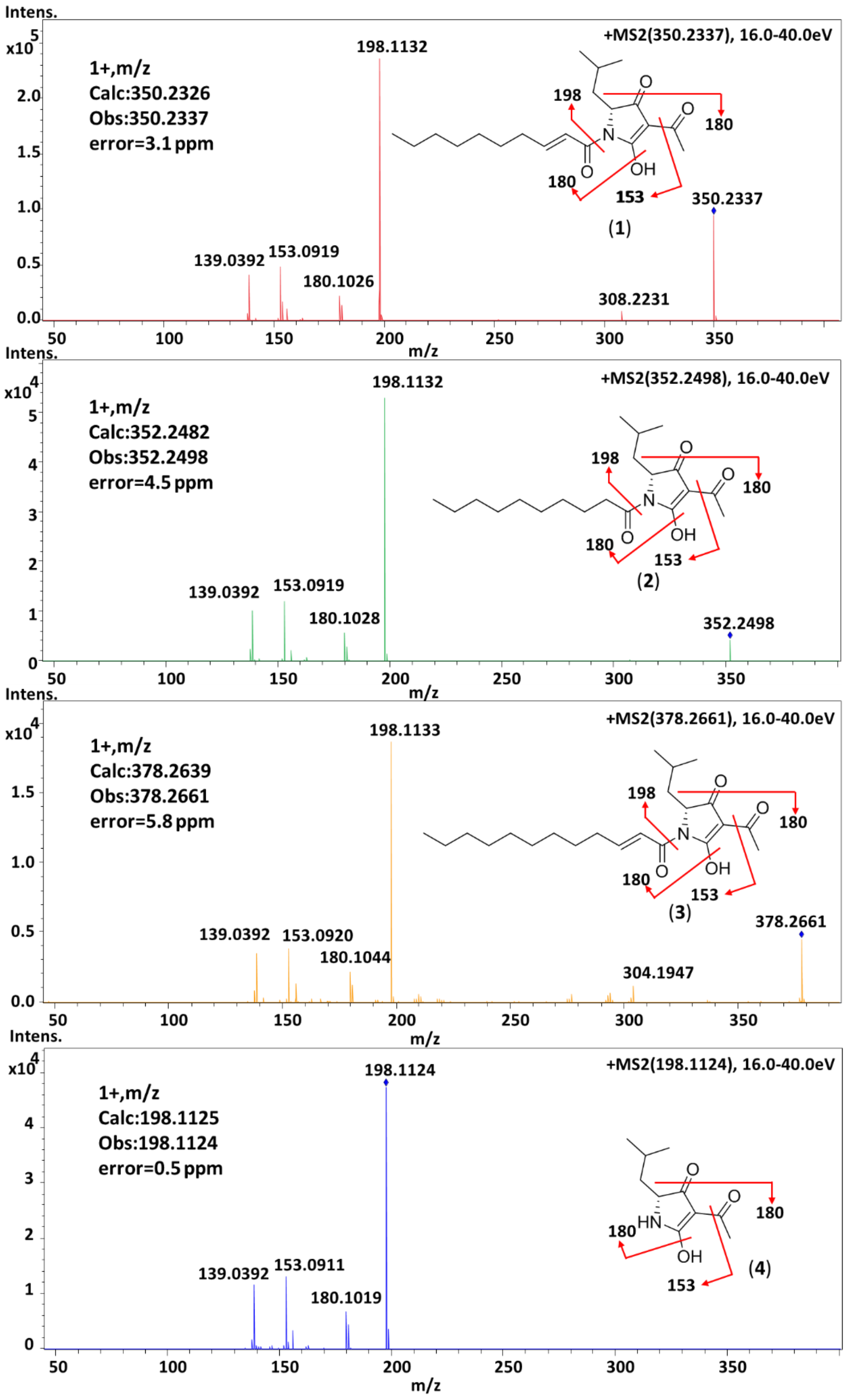

**Supplementary Figure 3.** $^1$H NMR spectrum of **1**.

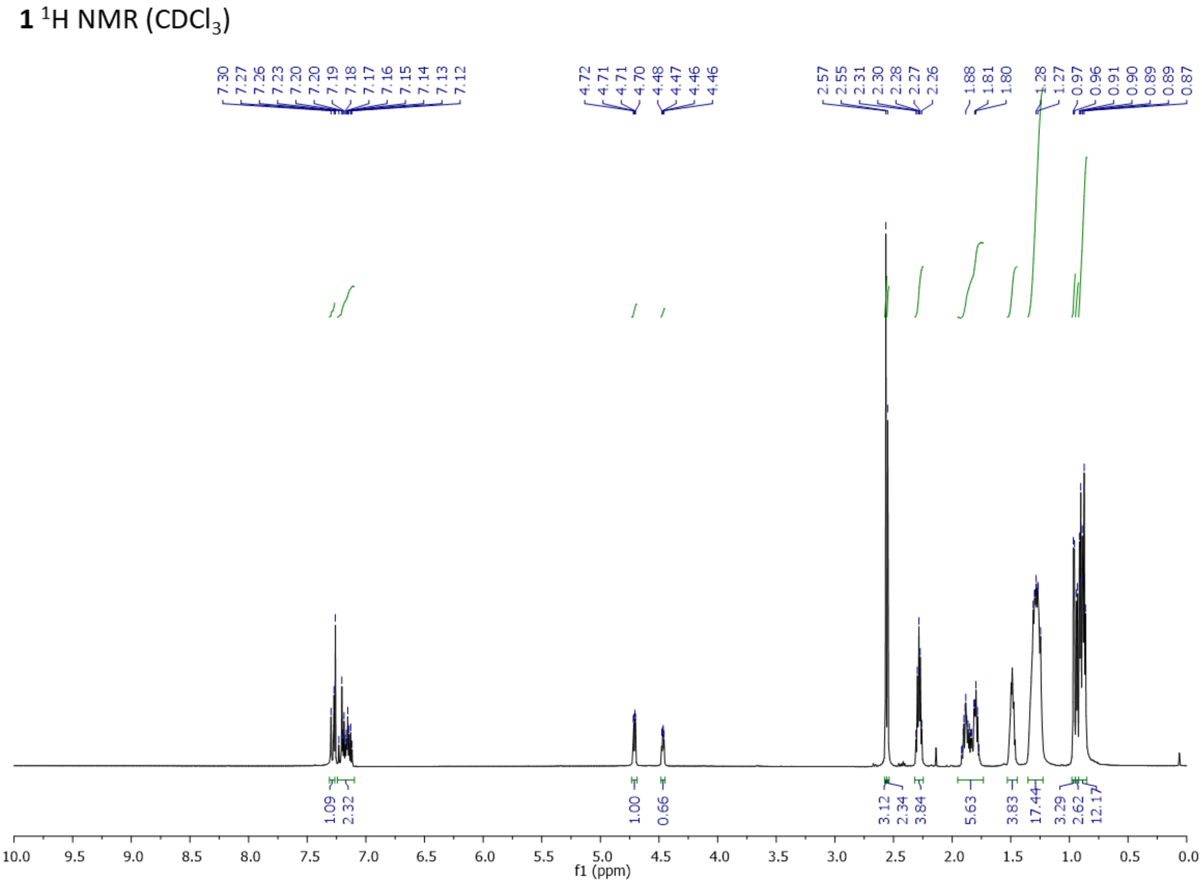

**Supplementary Figure 4.** $^{13}$C NMR spectrum of **1**.

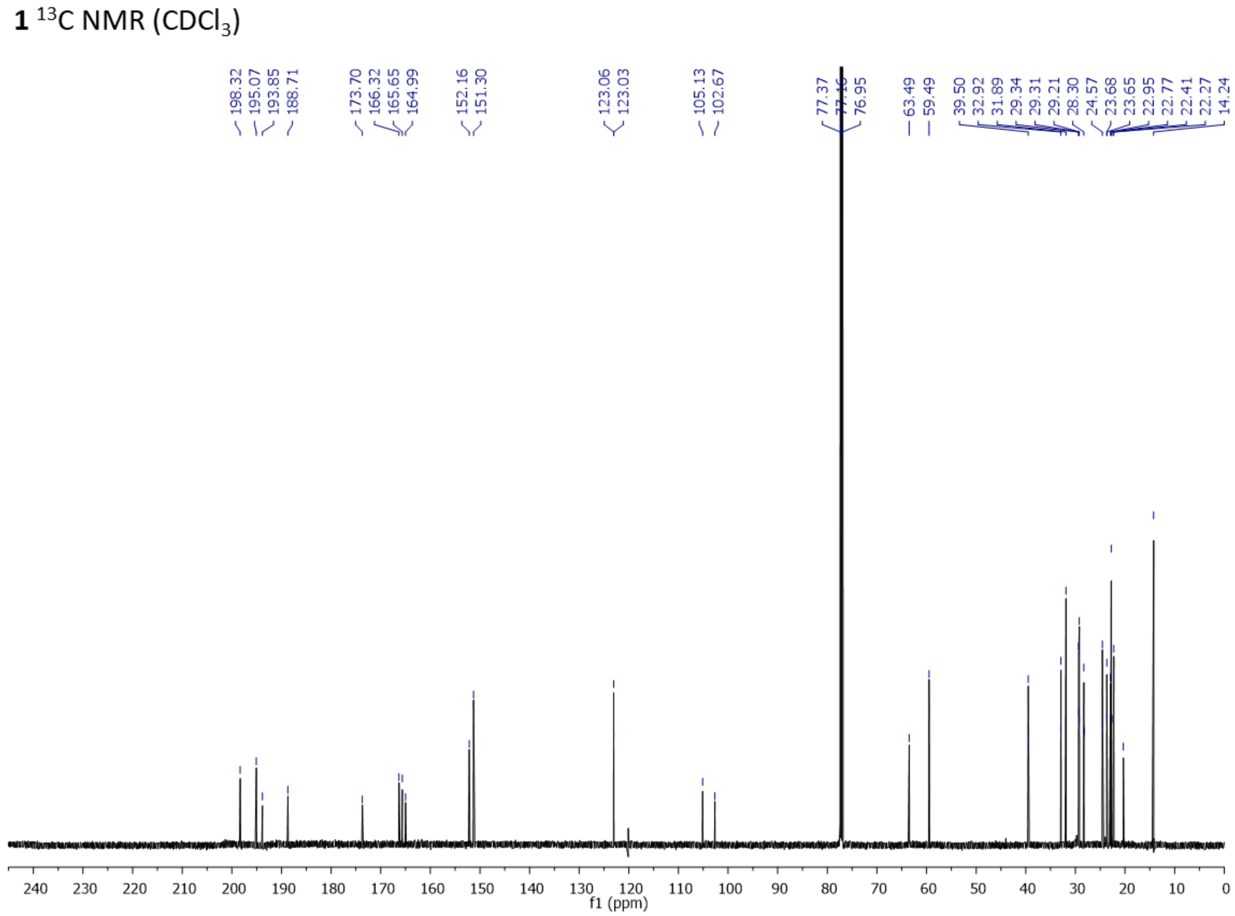

**Supplementary Figure 5.** COSY spectrum of **1**.

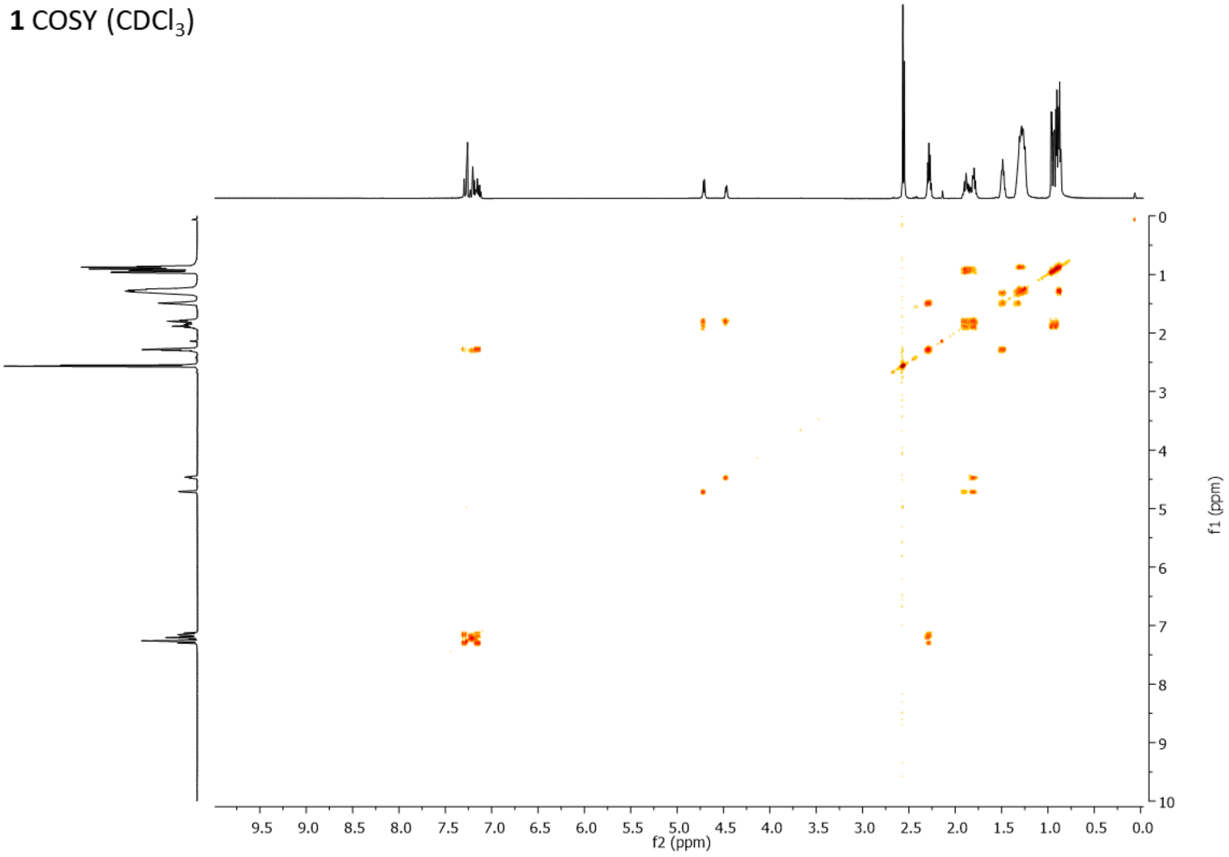

**Supplementary Figure 6.** HSQC spectrum of **1**.

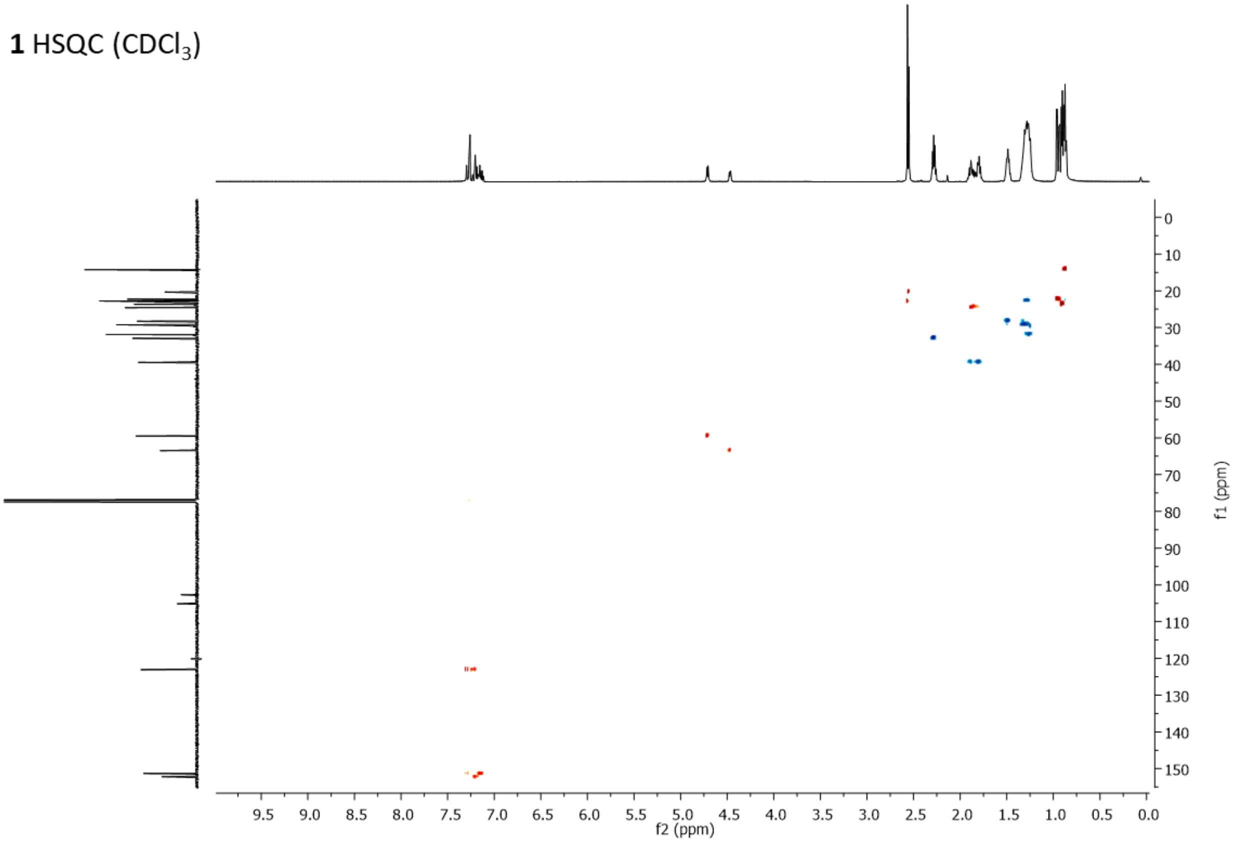

**Supplementary Figure 7.** HMBC spectrum of **1**.

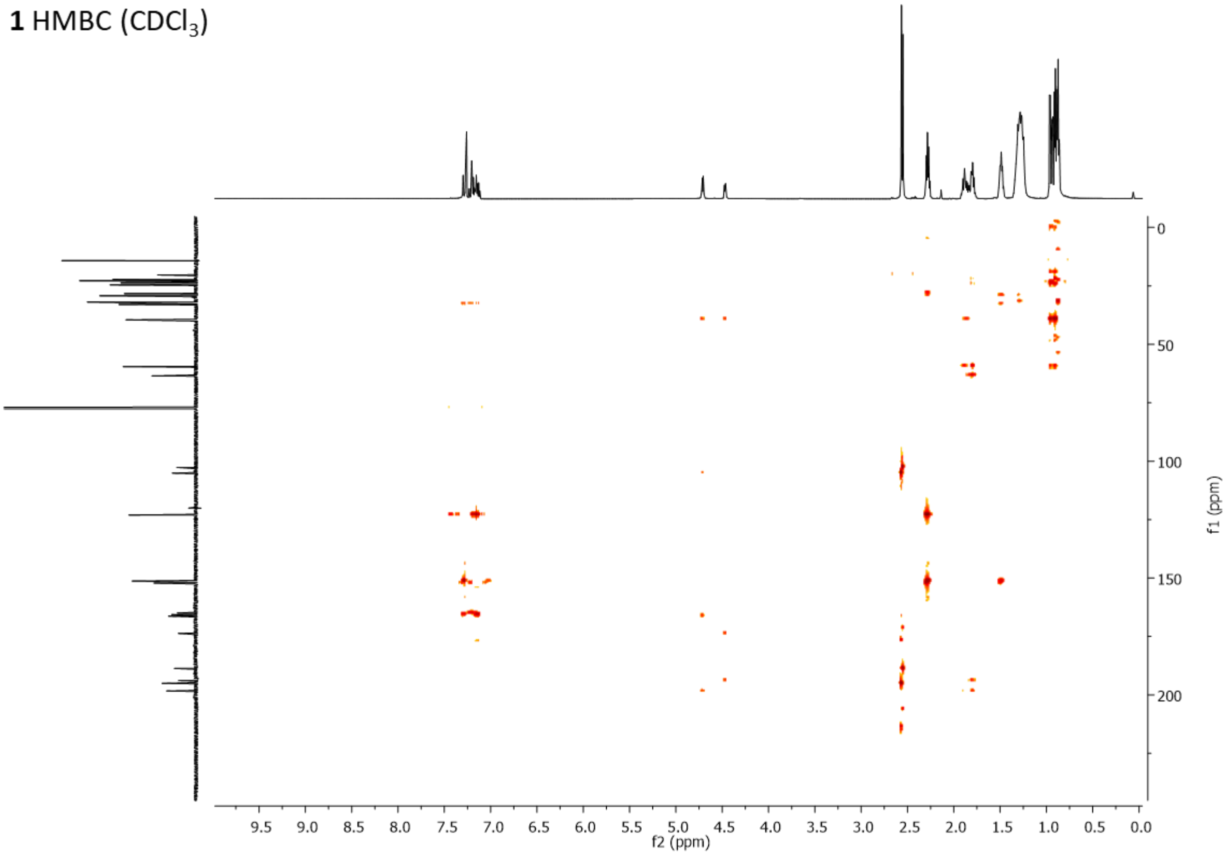

**Supplementary Figure 8.** Comparison of the isolated **1** with synthetic standards using (a) $^{1}H$ NMR, (b) $^{13}C$ NMR.

(a)

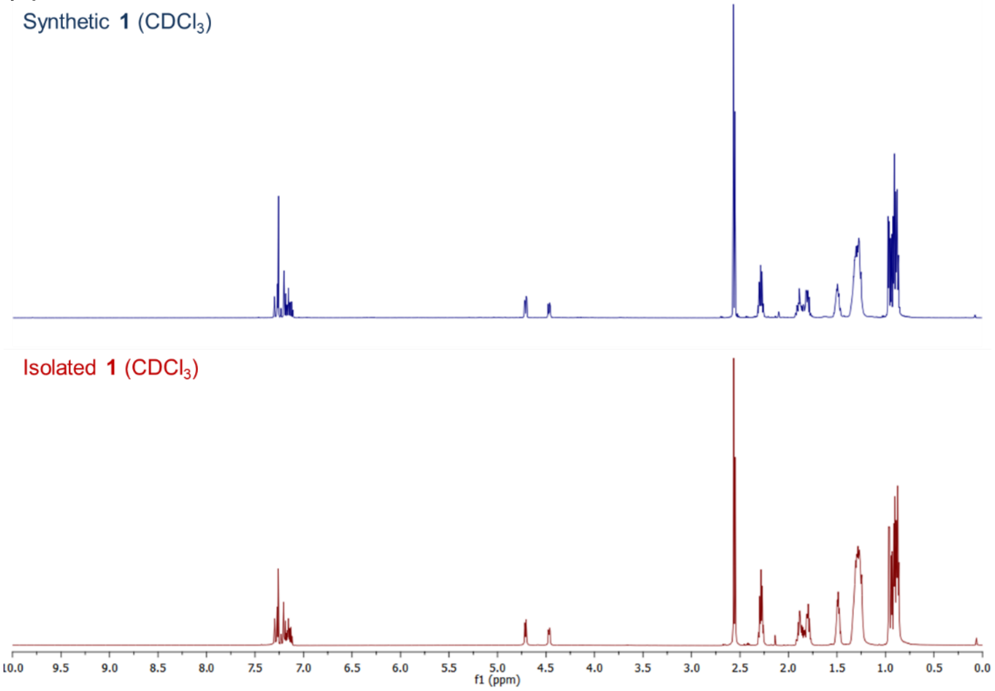

(b)

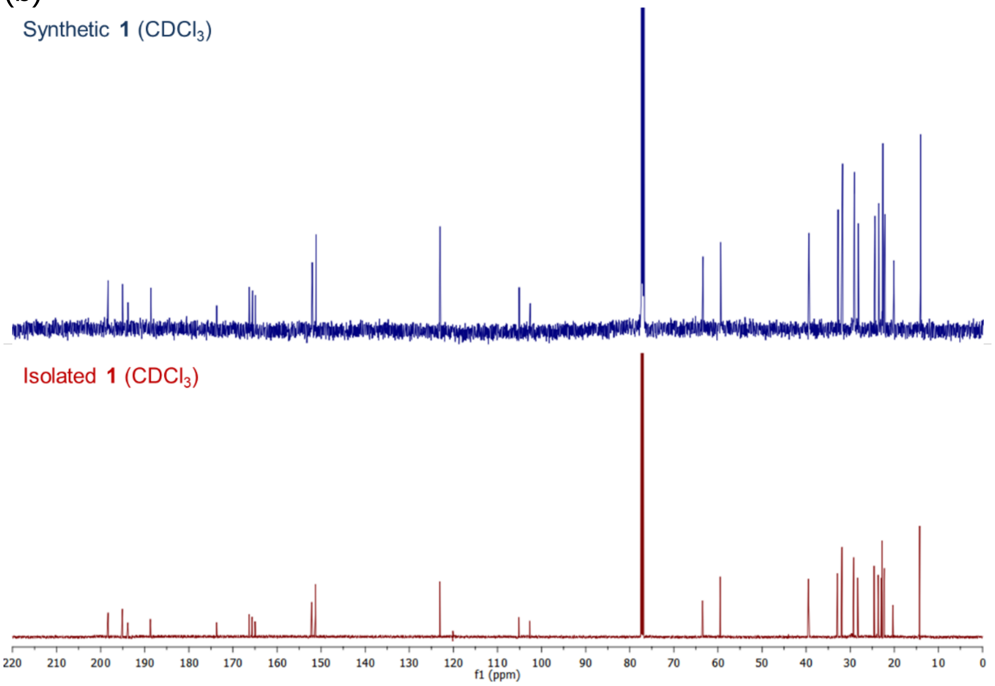

**Supplementary Figure 9.** Comparison of compound **1** with synthetic and isolated standards by chiral HPLC. (a) Synthetic standard was derived from (*S*)-**4** (22.7% ee). (b) **1** isolated from *S. mutans*. (c) **1** isolated from *Lactobacillus reuteri* LTH2584(1).

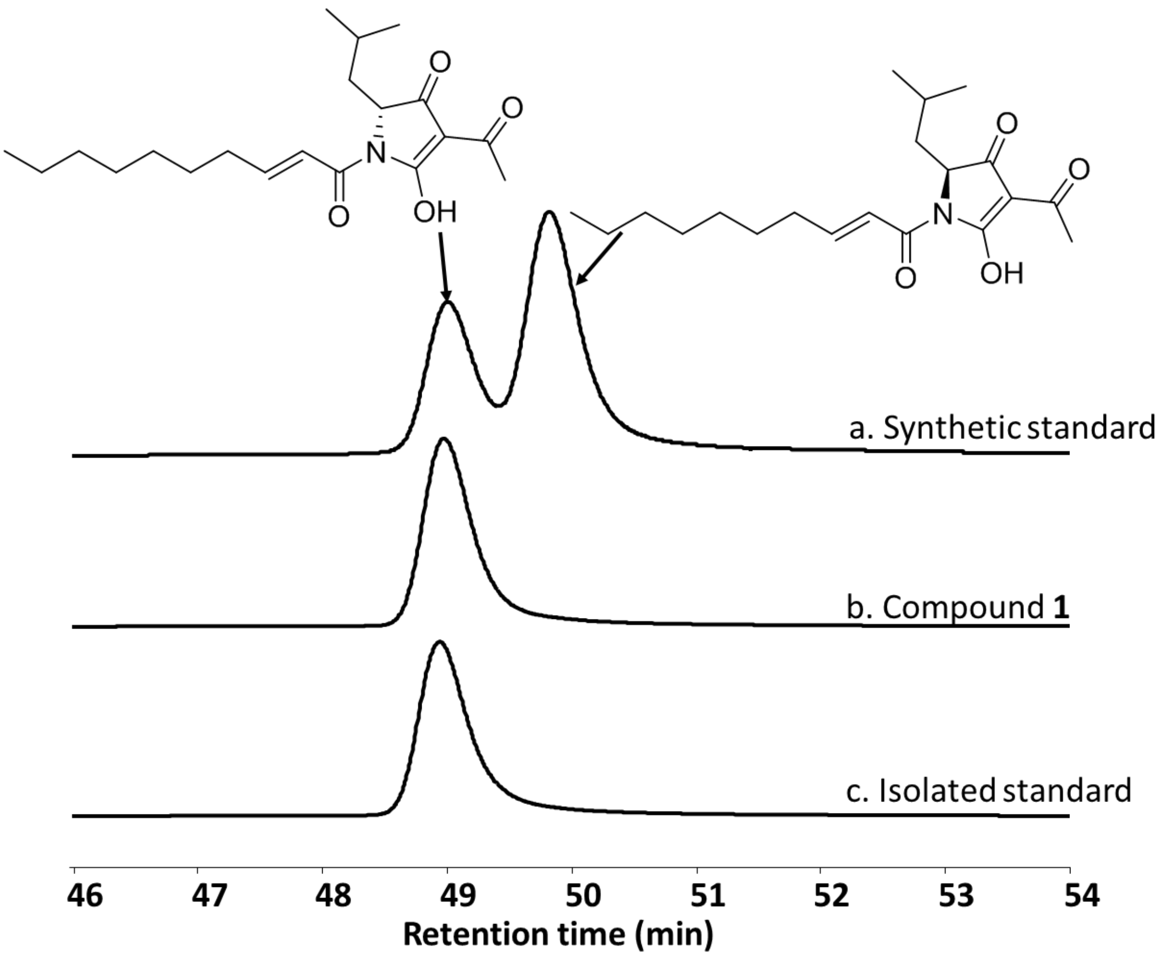

**Supplementary Figure 10.** $^{1}H$ NMR spectrum of **3**.

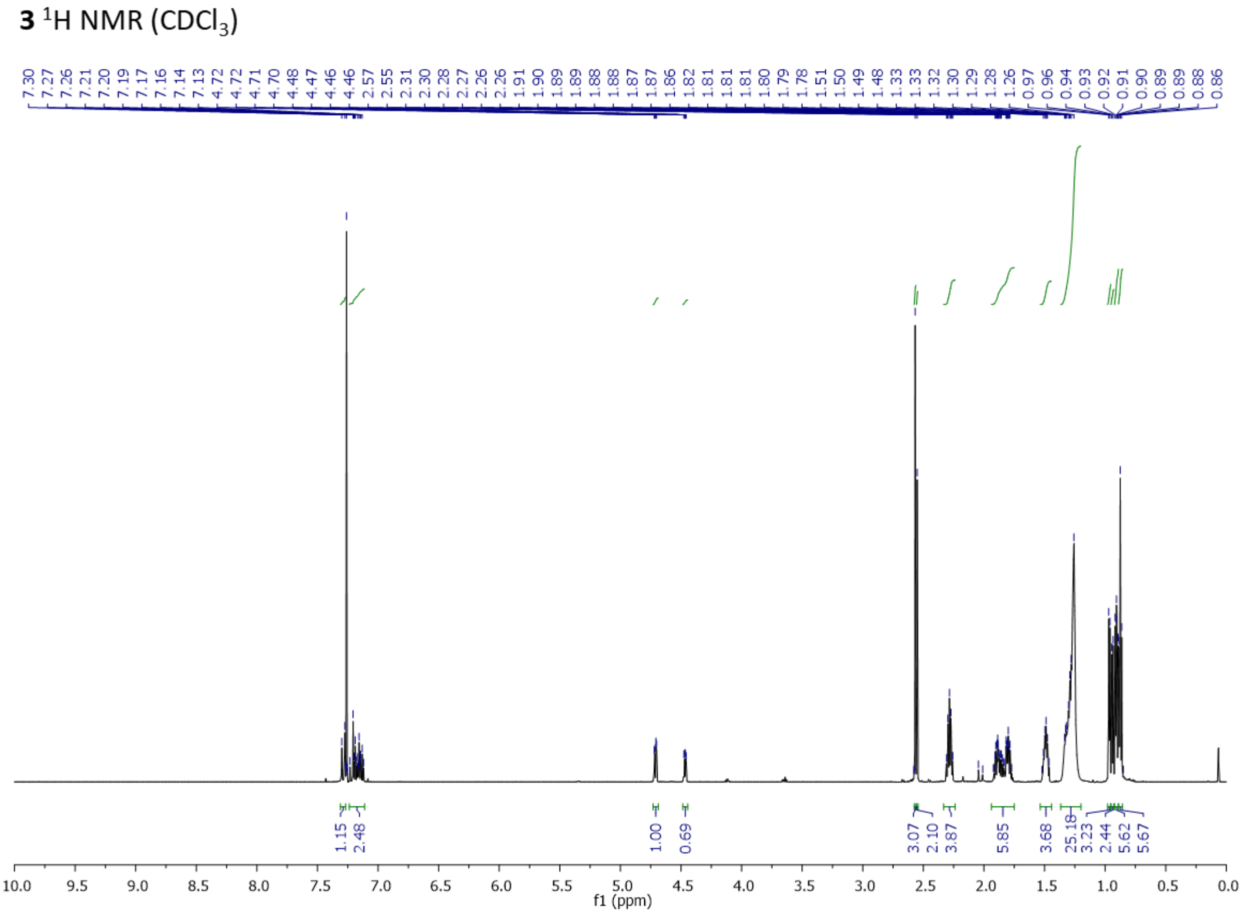

**Supplementary Figure 11.** $^{13}$C NMR spectrum of **3**.

**3** $^{13}$C NMR ($CDCl_3$)

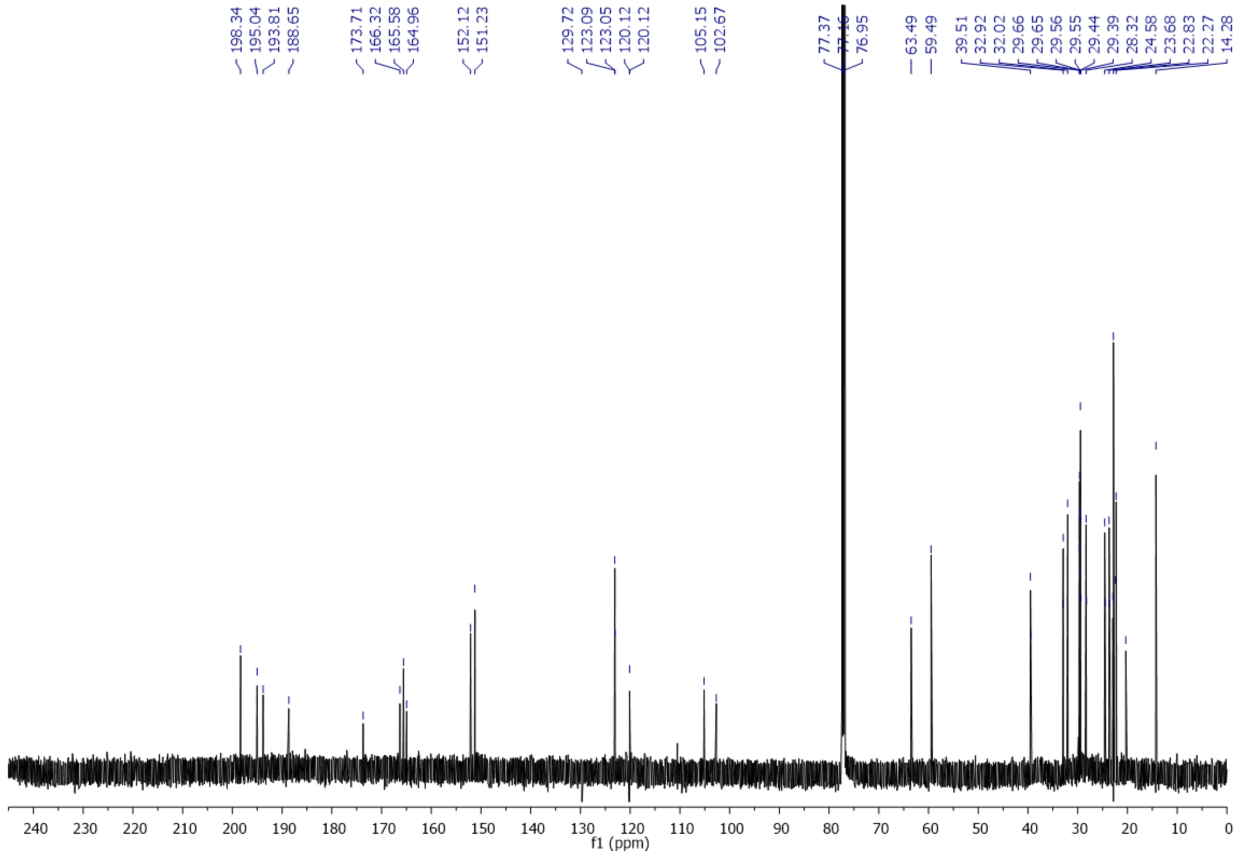

**Supplementary Figure 12**. COSY spectrum of **3**.

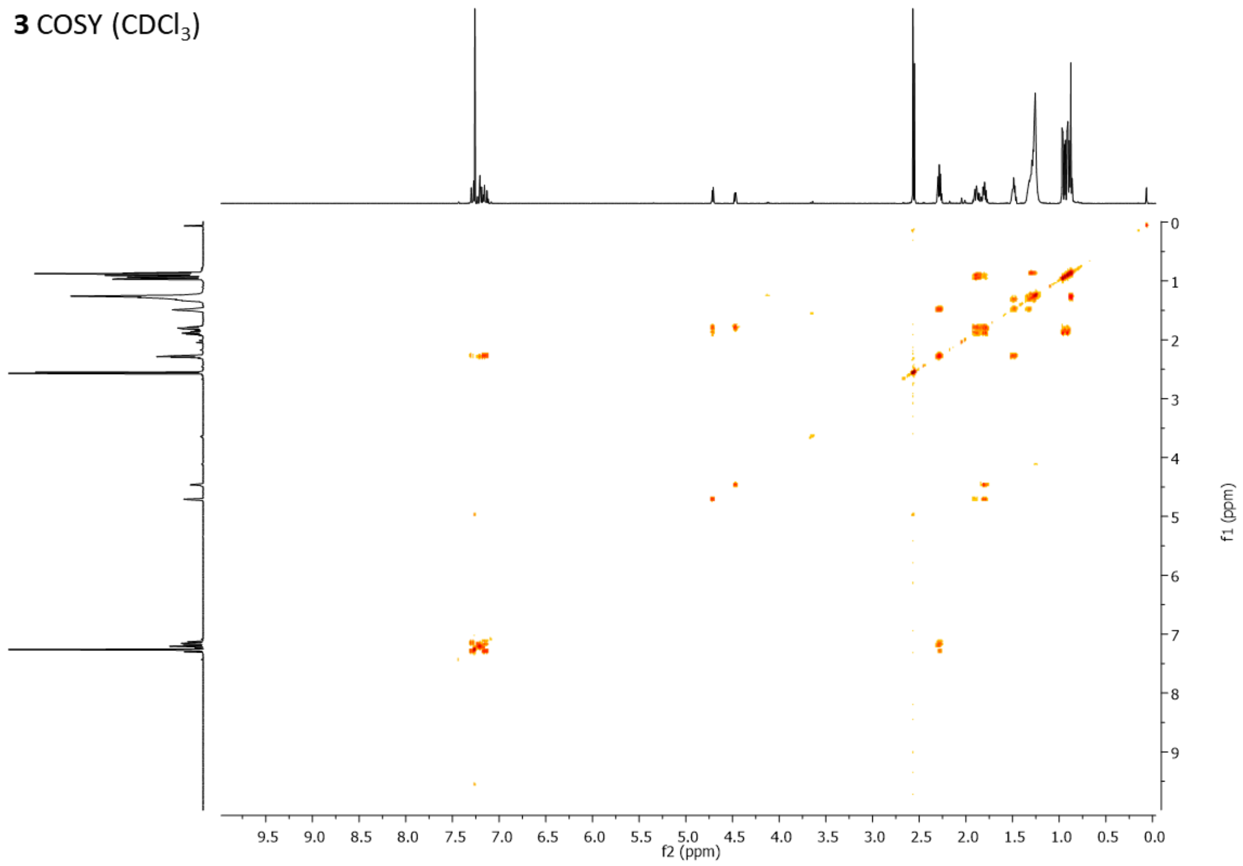

**Supplementary Figure 13**. HSQC spectrum of **3**.

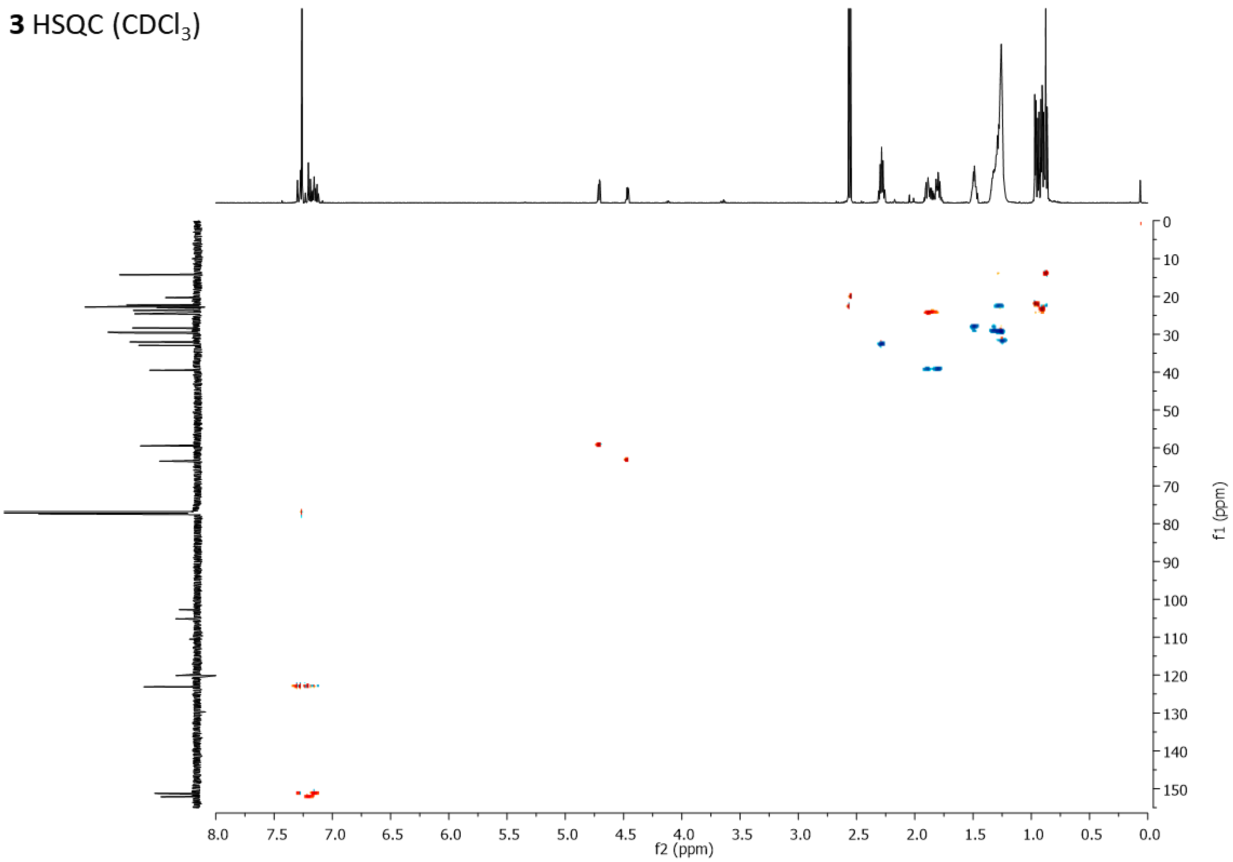

**Supplementary Figure 14**. HMBC spectrum of **3**.

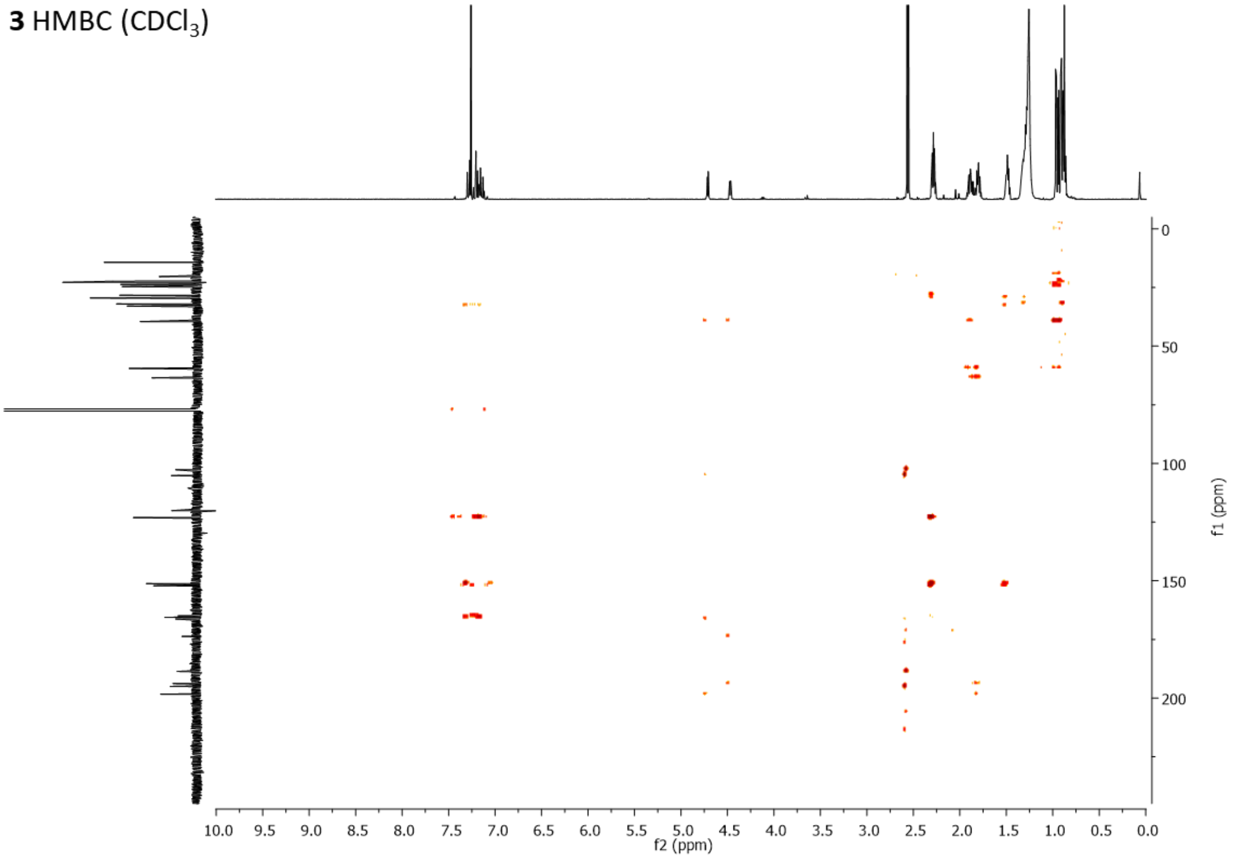

**Supplementary Figure 15.** Comparison of the isolated **4** with synthetic standards using (a) HPLC, (b) LC-MS/MS, and (c) $^{1}$H-NMR.

**(a)**

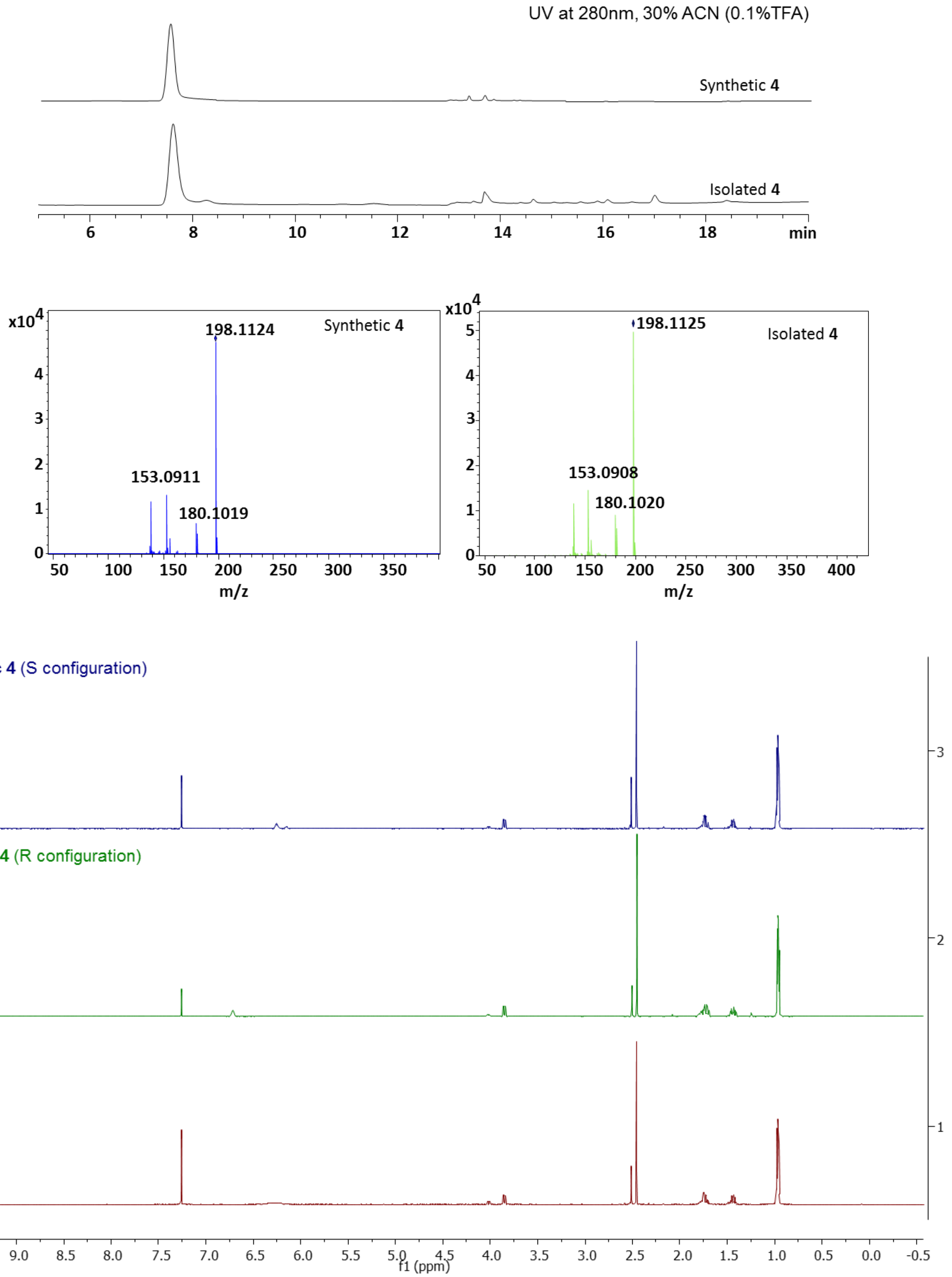

**Supplementary Figure 16.** Comparison of **4** with synthetic standards by chiral HPLC. (a) Compound **4** isolated from *S. mutans*. (b) Purified synthetic (*R*)-**4** standard; (c) Purified synthetic (*S*)-**4** standard.

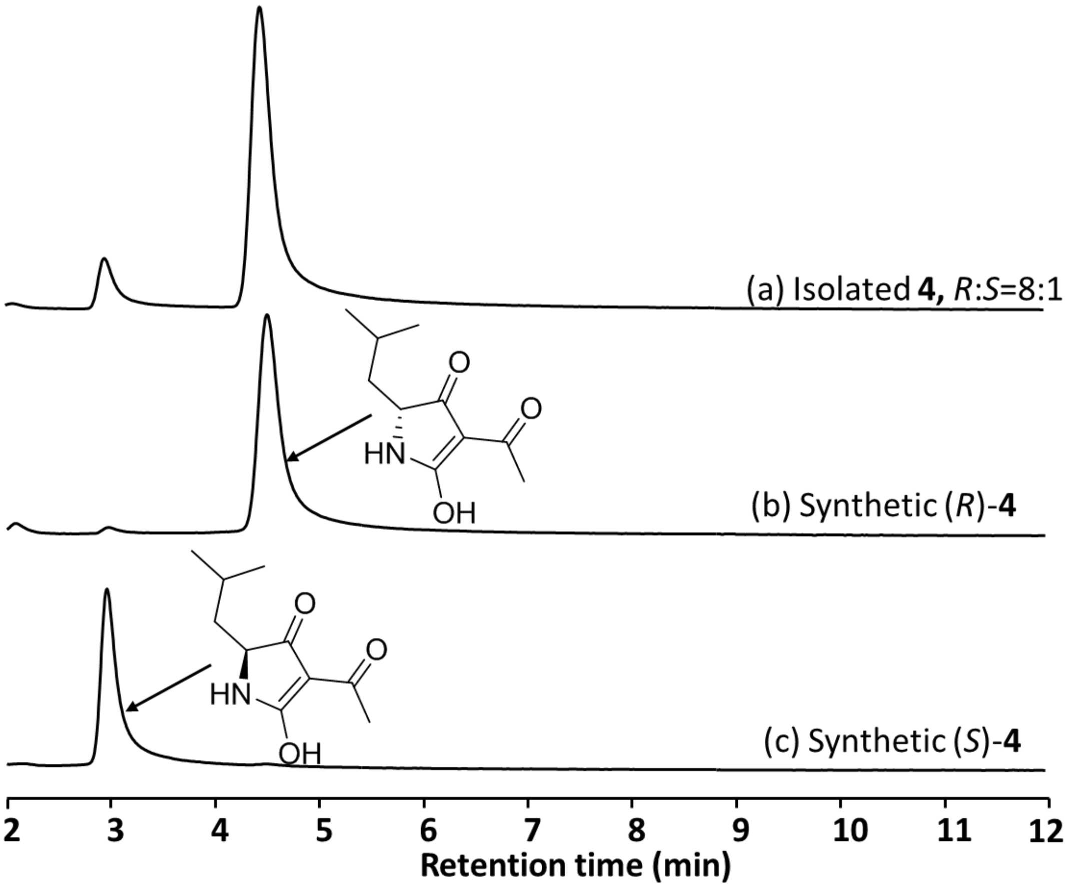

**Supplementary Figure 17.** $^{1}$H NMR spectrum of synthetic (*R*)-**4**.

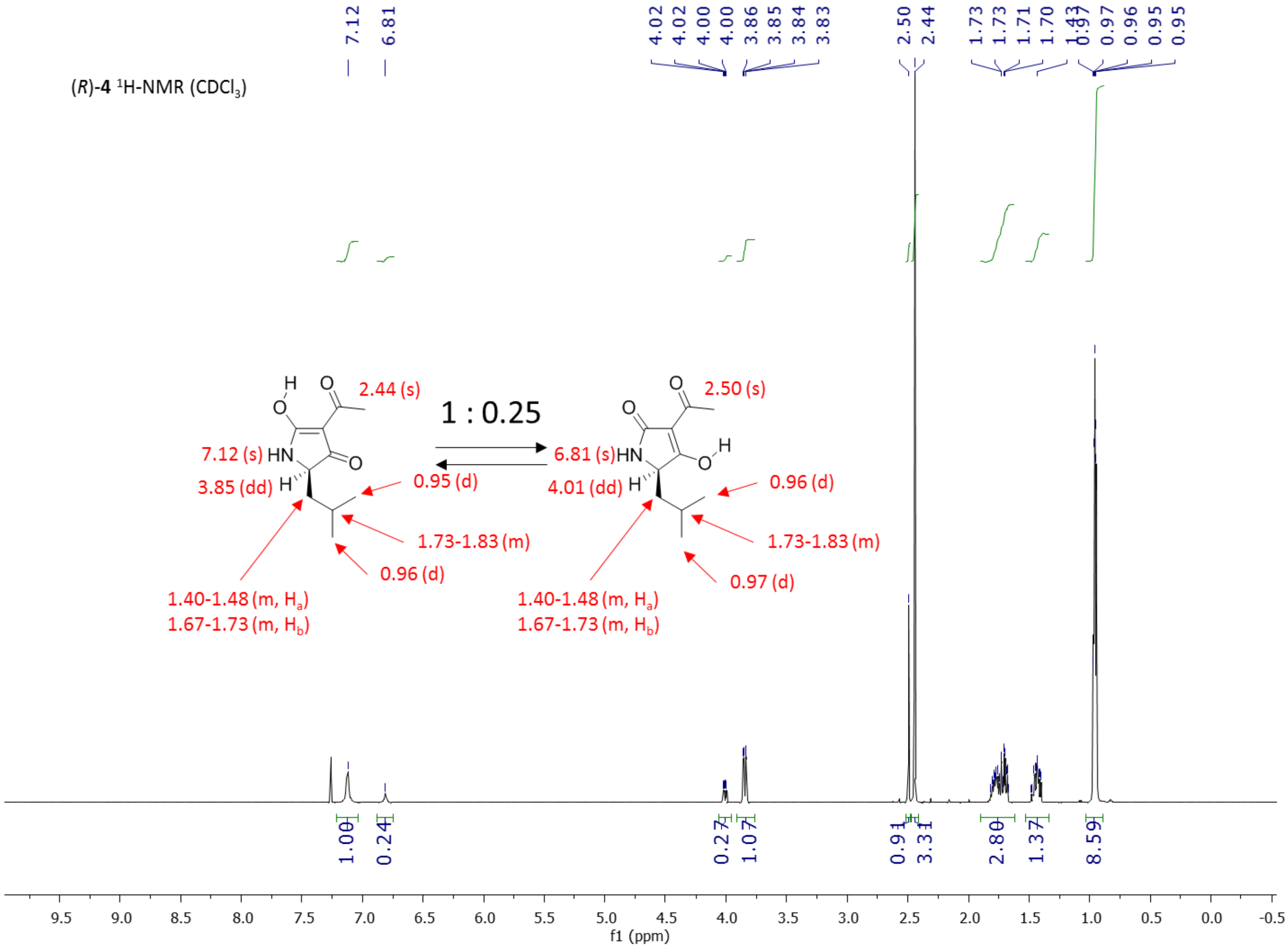

**Supplementary Figure 18**. $^{13}$C NMR spectrum of synthetic (*R*)-**4**.

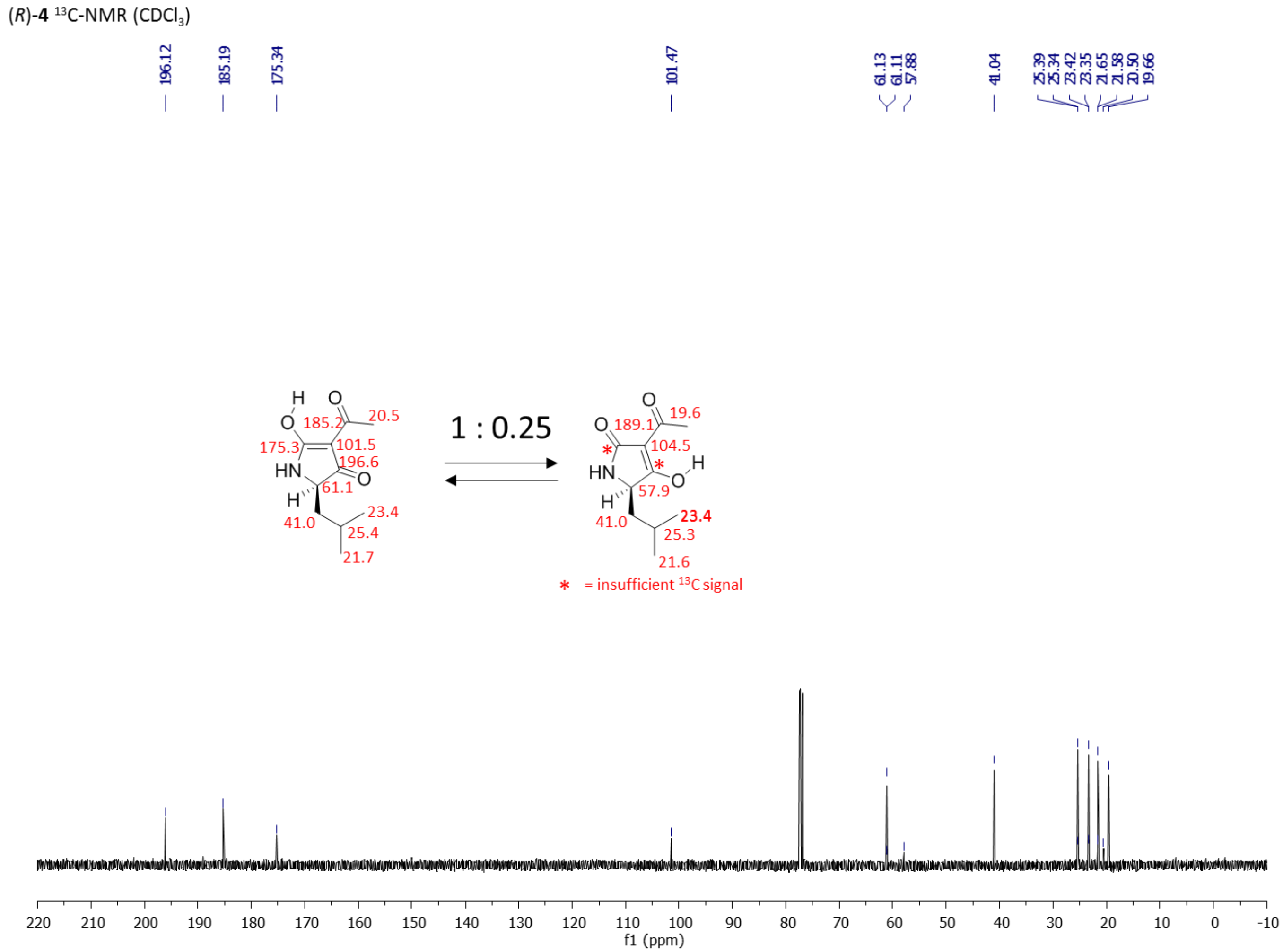

**Supplementary Figure 19.** COSY spectrum of synthetic (*R*)-**4**.

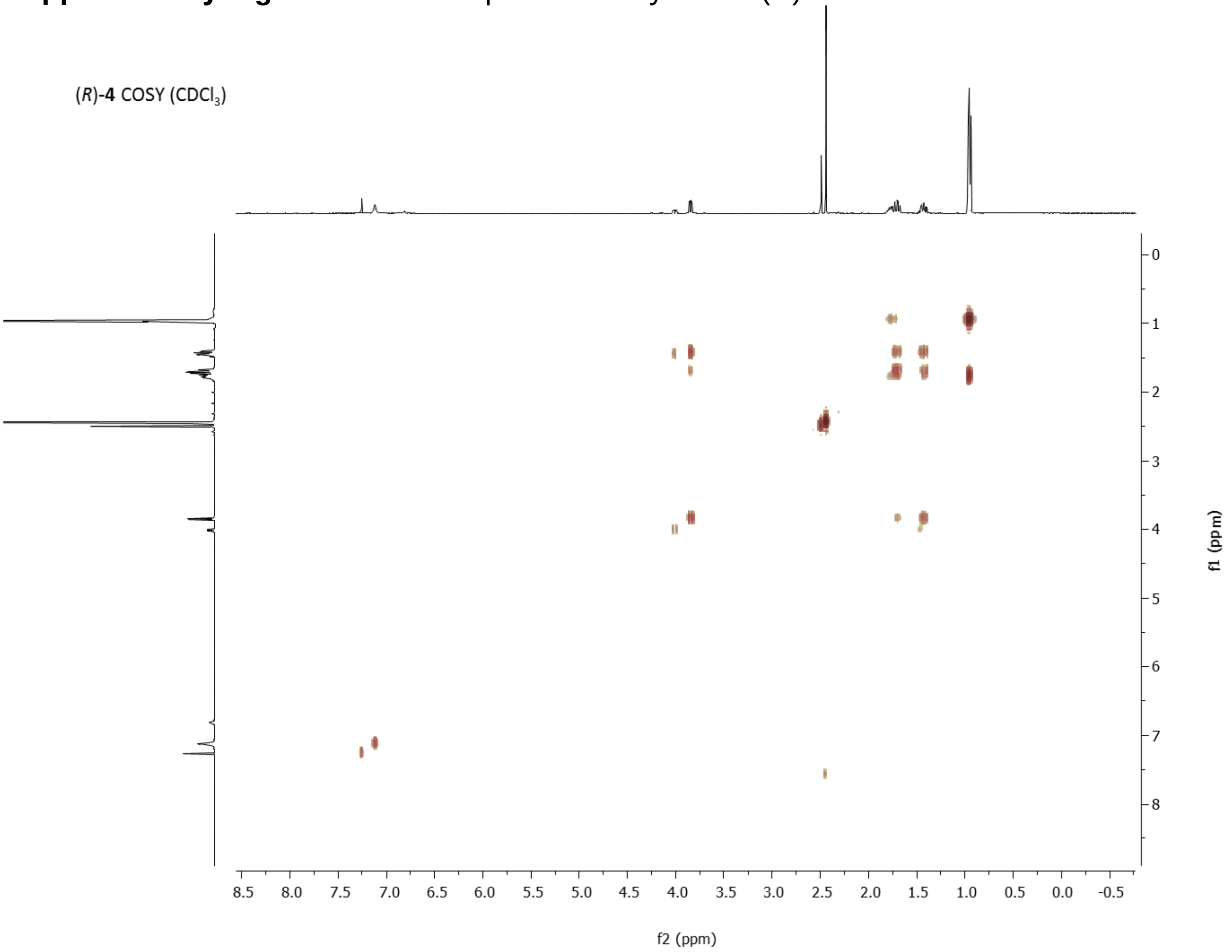

**Supplementary Figure 20.** HSQC spectrum of synthetic (*R*)-**4**.

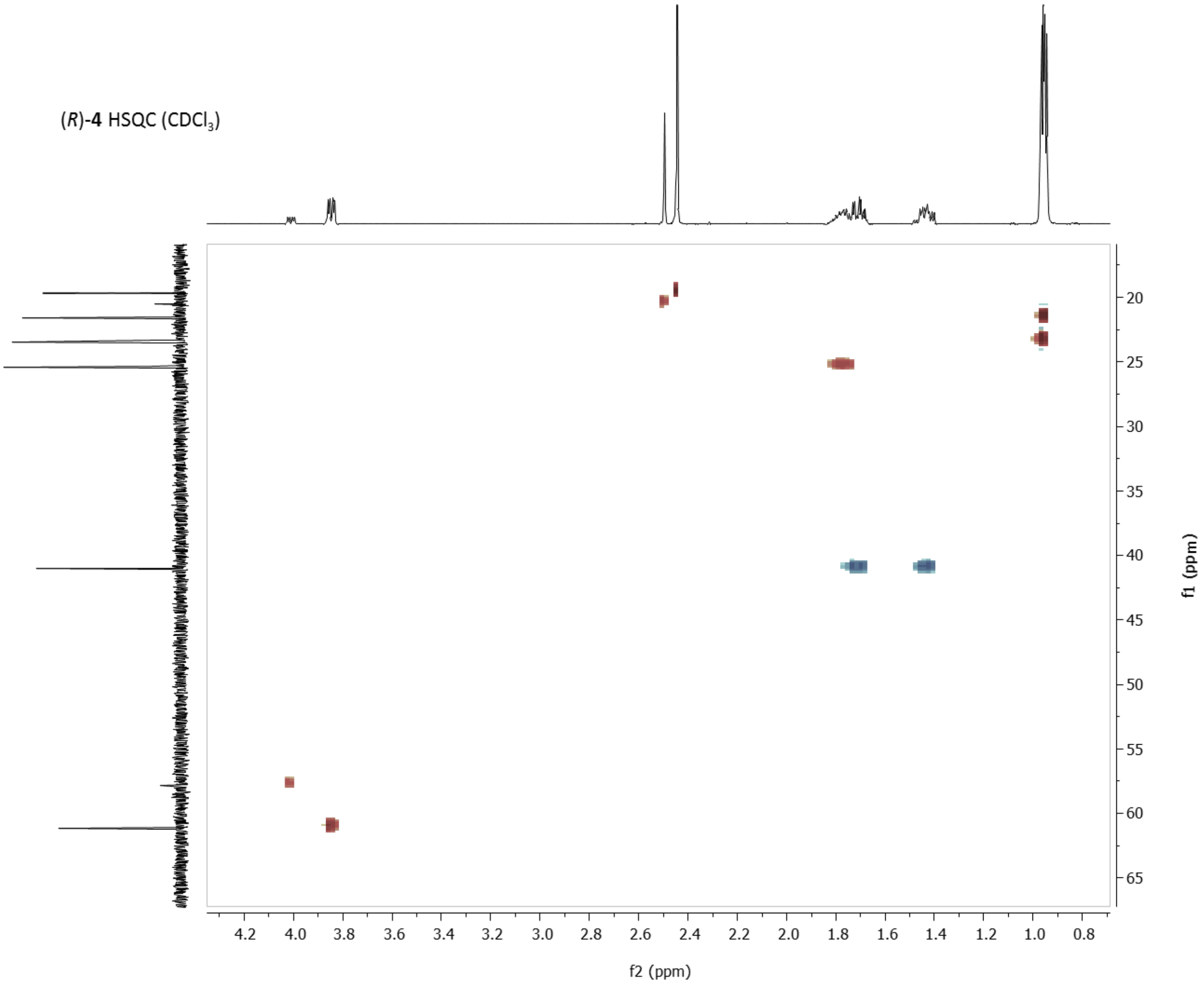

**Supplementary Figure 21.** HMBC spectrum of synthetic (*R*)-**4**.

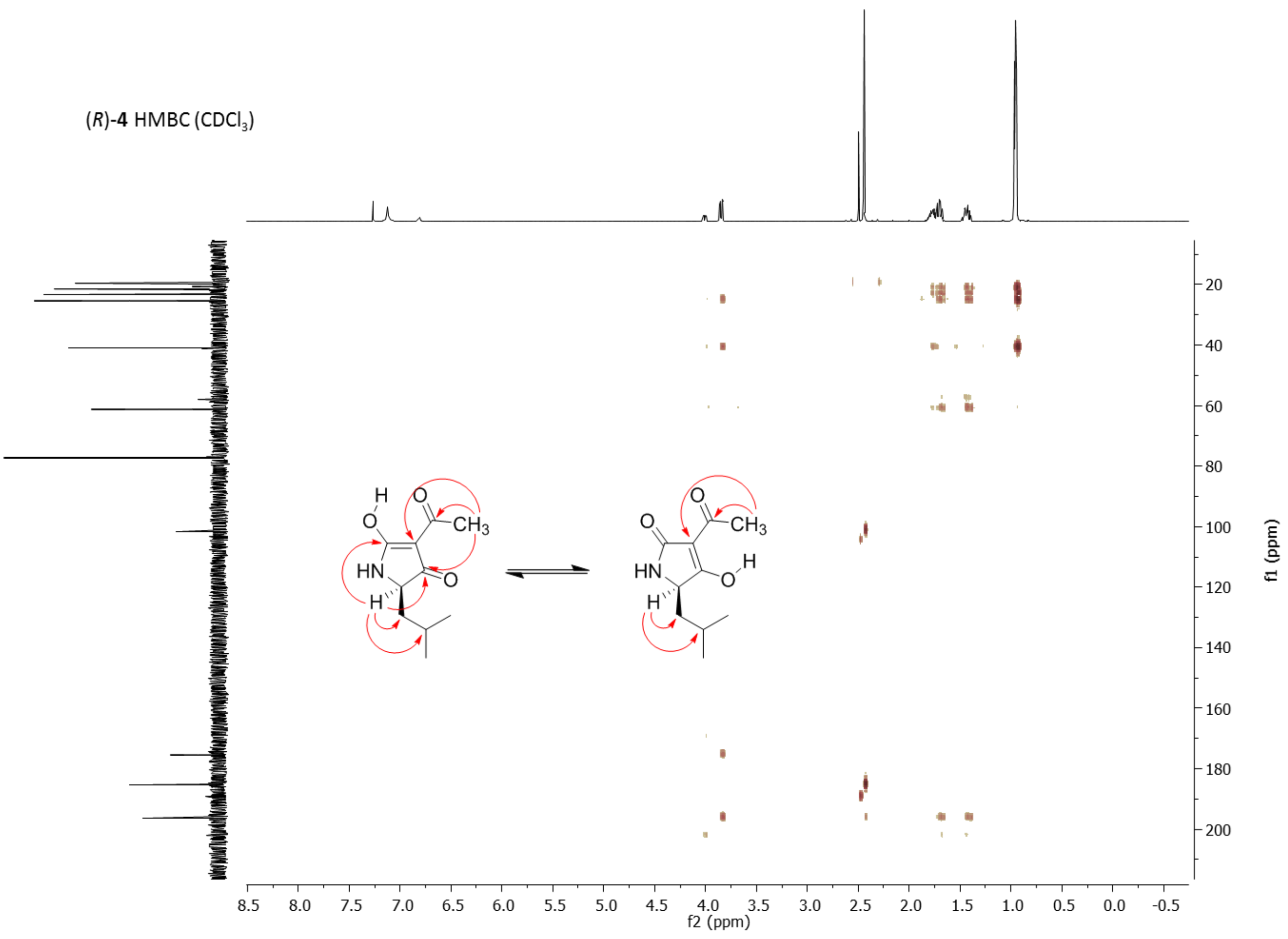

**Supplementary Figure 22.** Chiral HPLC analysis of synthetic (*S*)-**4** and (*R*)-**4**.

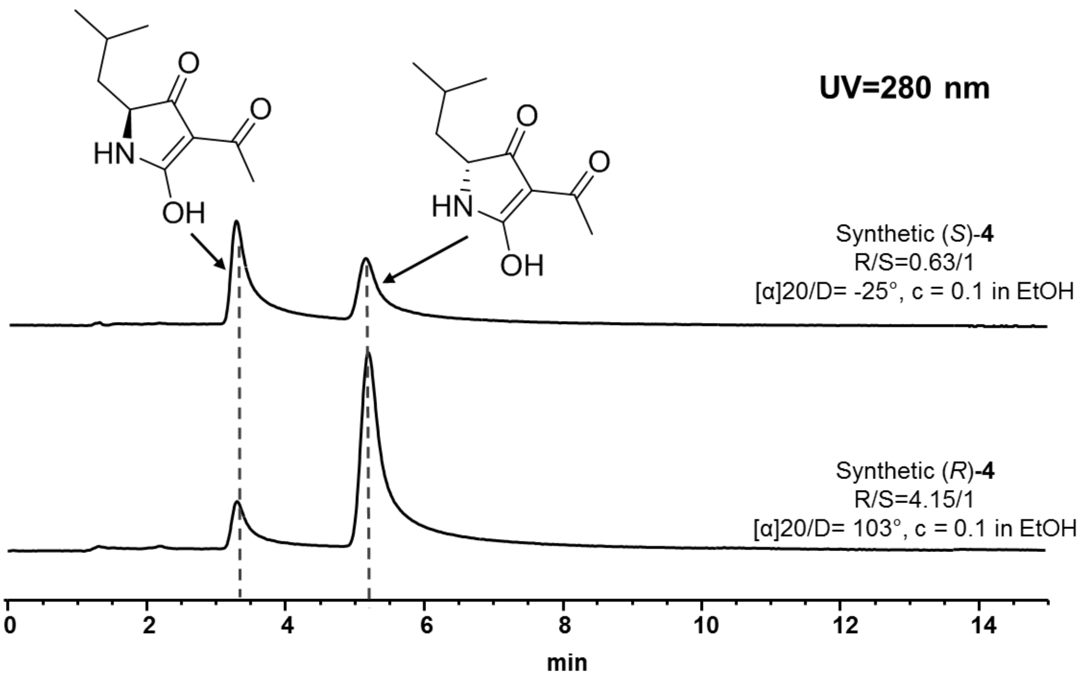

**Supplementary Figure 23.** MS analysis of **1** isolated from *S. mutans* B04Sm5 feeding experiments.

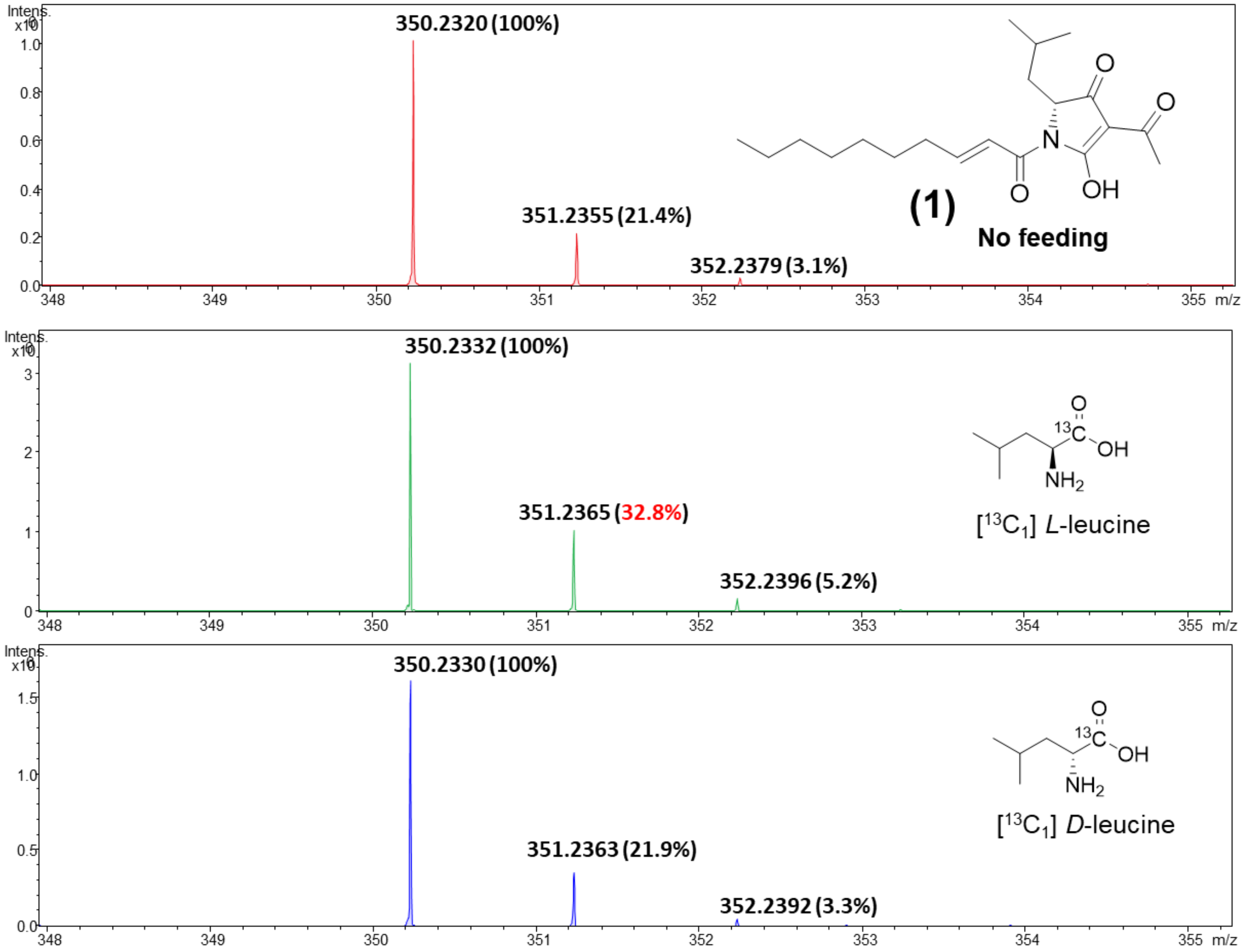

**Supplementary Figure 24.** MS analysis of **3** isolated from *S. mutans* B04Sm5 feeding experiments.

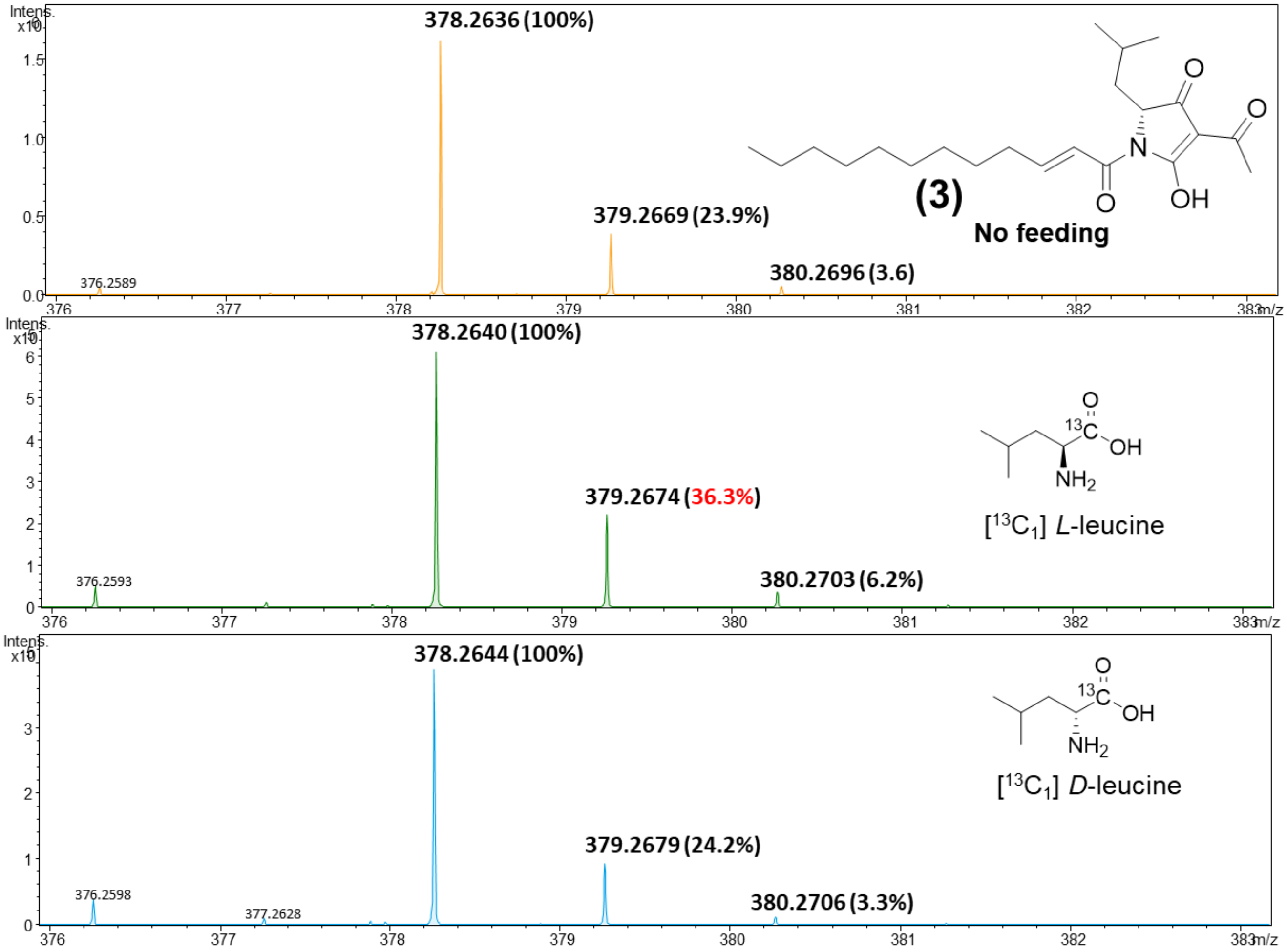

**Supplementary Figure 25.** MS analysis of **1** isolated from *L. reuteri* LTH2584(1) feeding experiments.

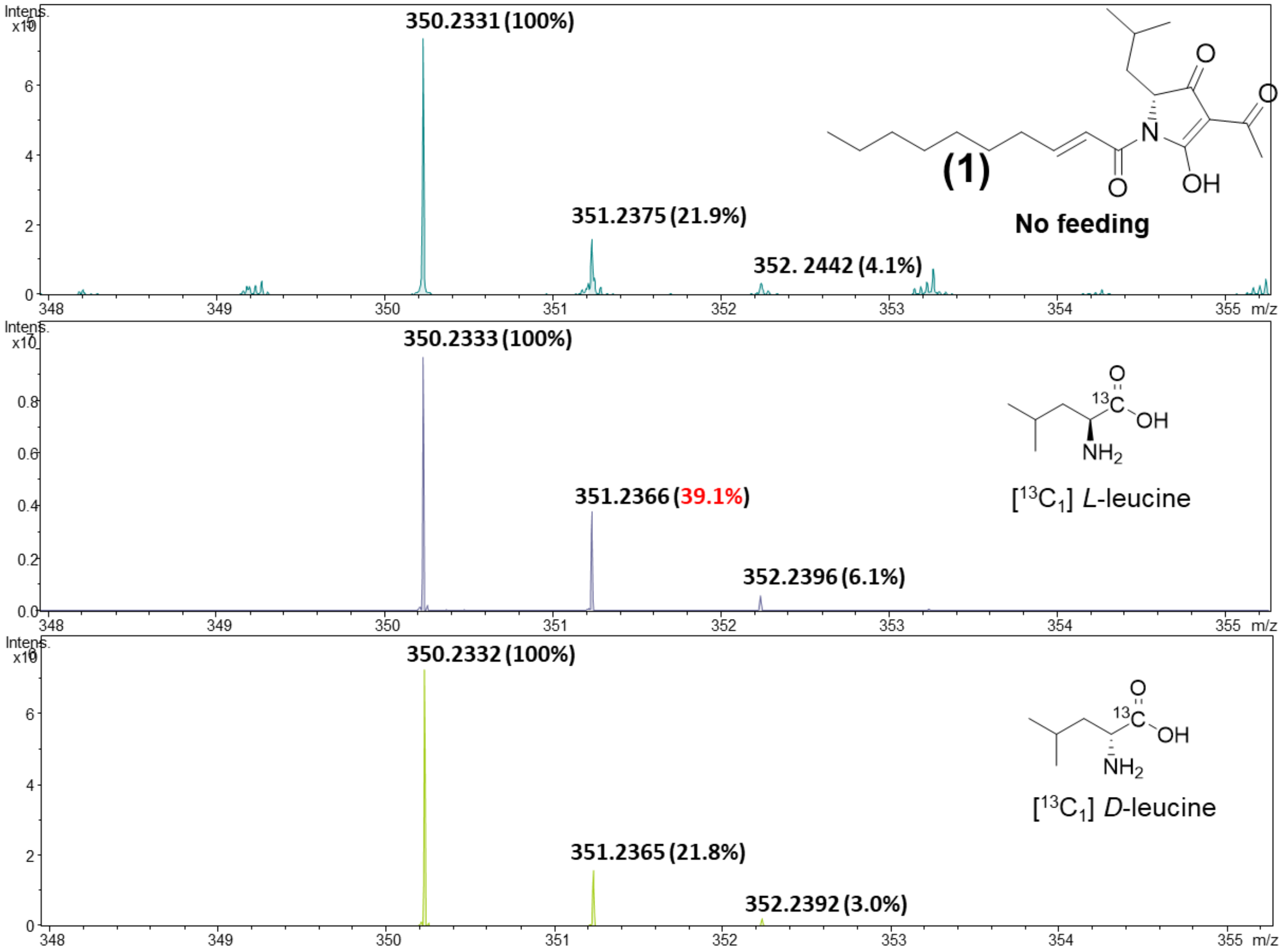

**Supplementary Figure 26**. High-resolution MS/MS spectrometry analysis of **1**-**5** isolated from *E. coli* BAP1 expression host.

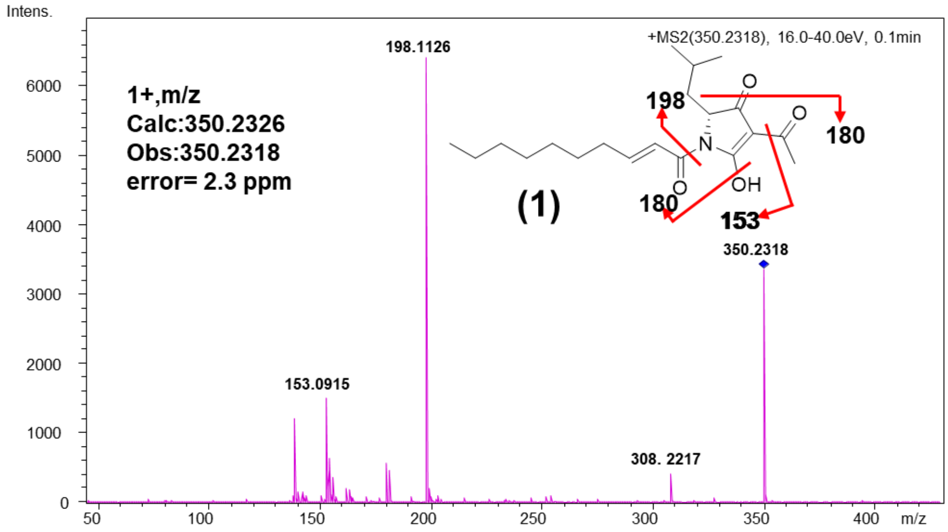

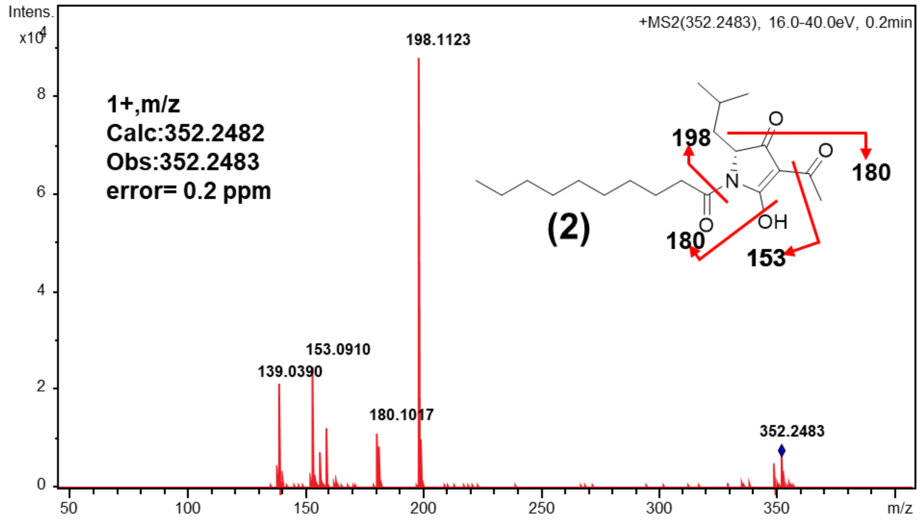

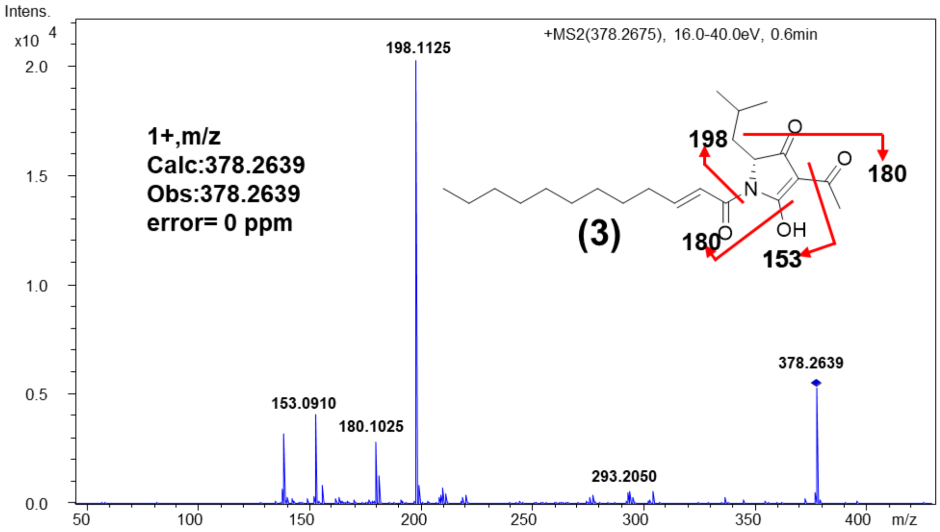

**Continued Supplementary Figure 26.**

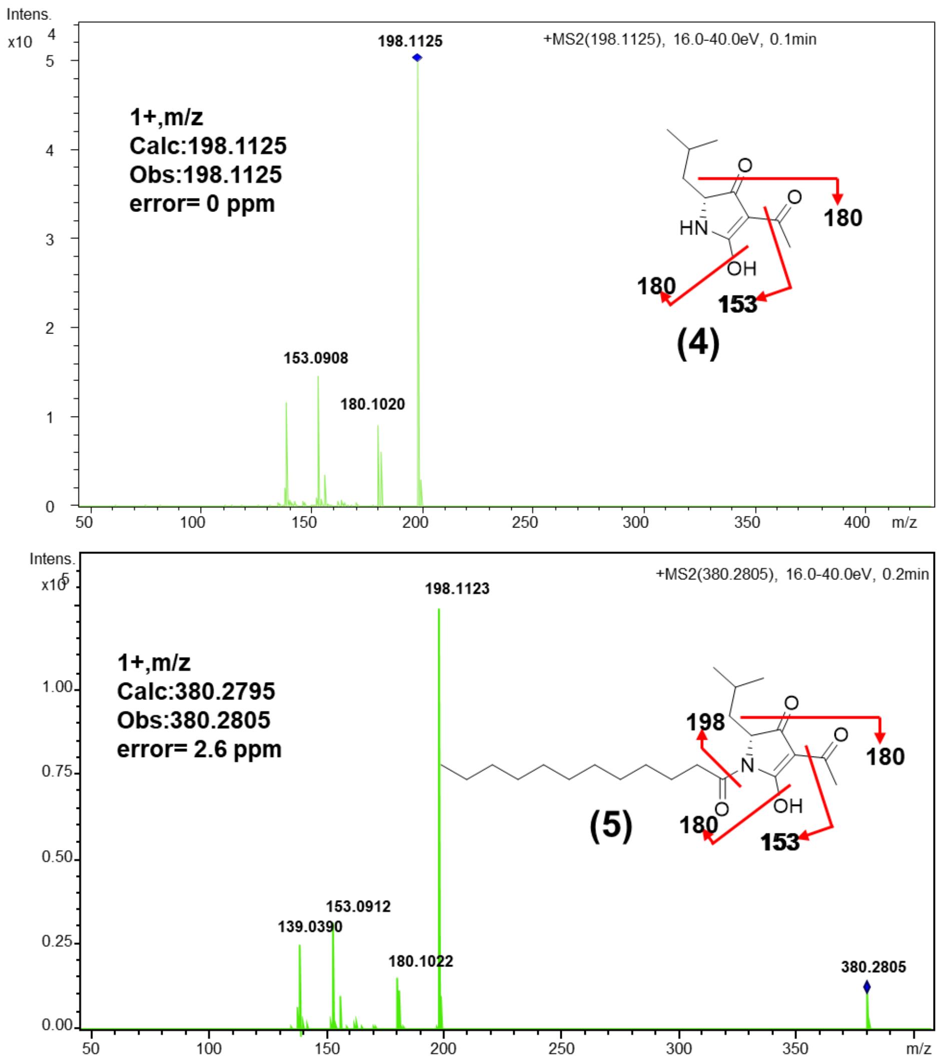

**Supplementary Figure 27.** $^{1}$H NMR spectrum of **5**.

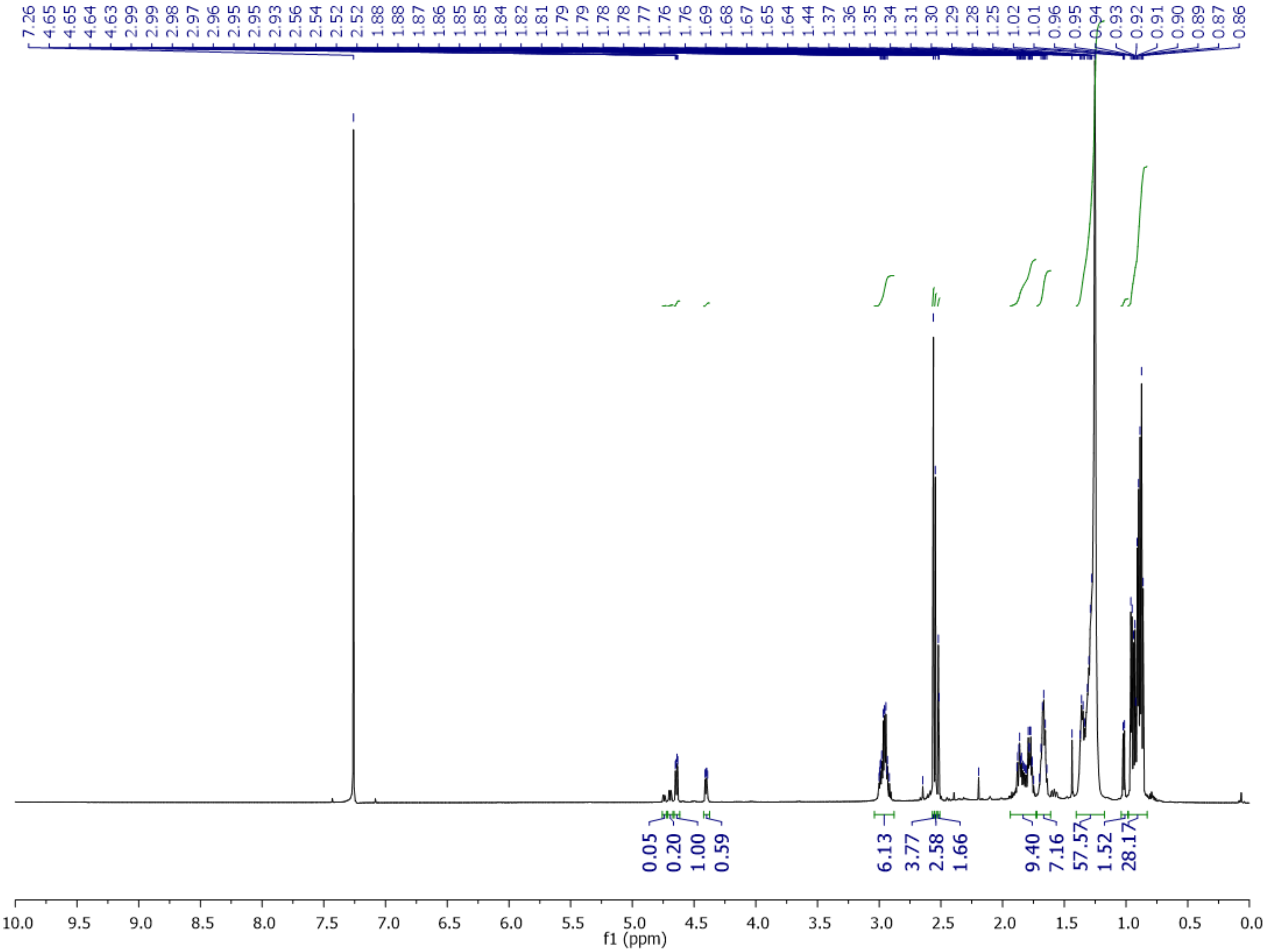

**Supplementary Figure 28.** COSY spectrum of **5**.

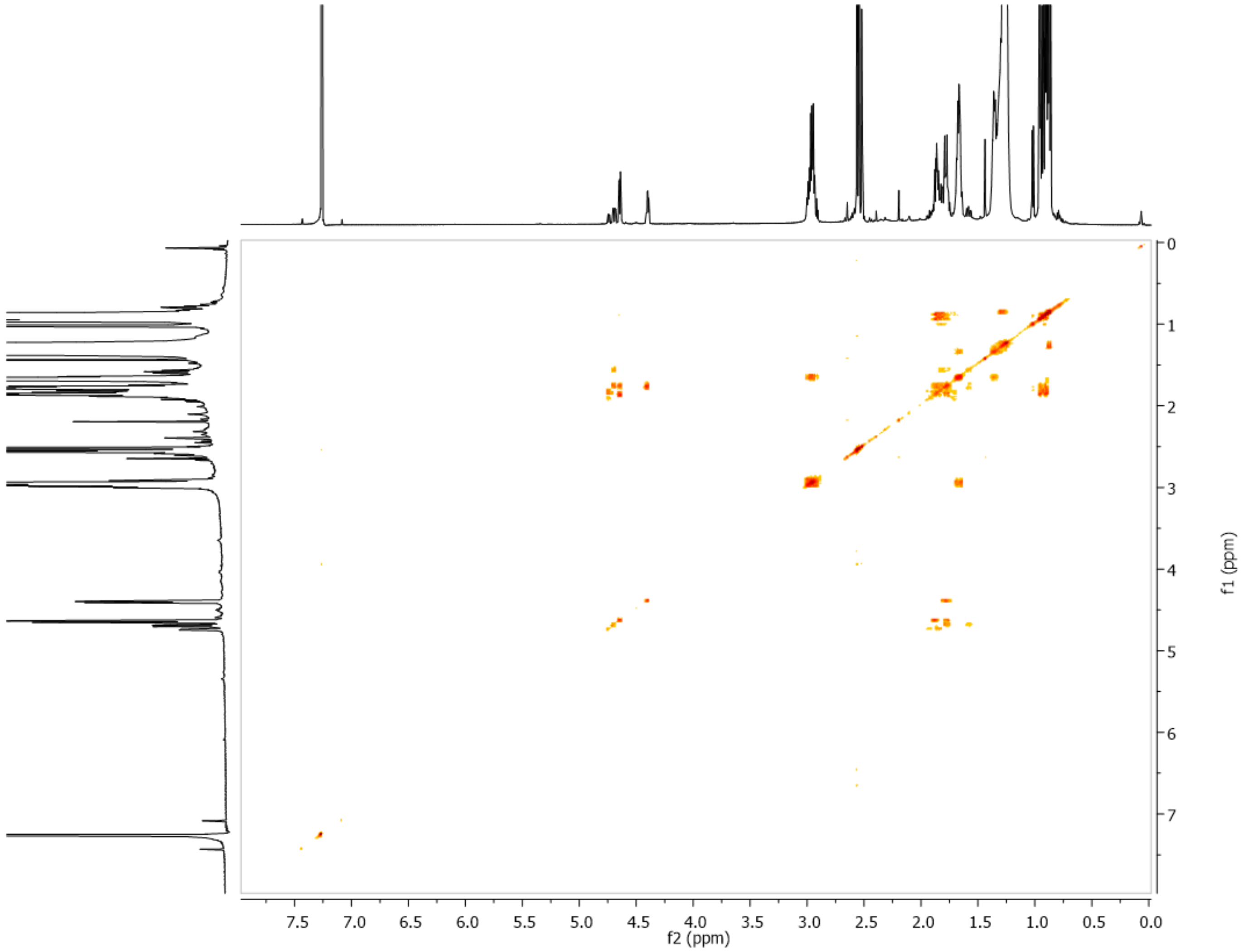

**Supplementary Figure 29.** HSQC spectrum of **5**

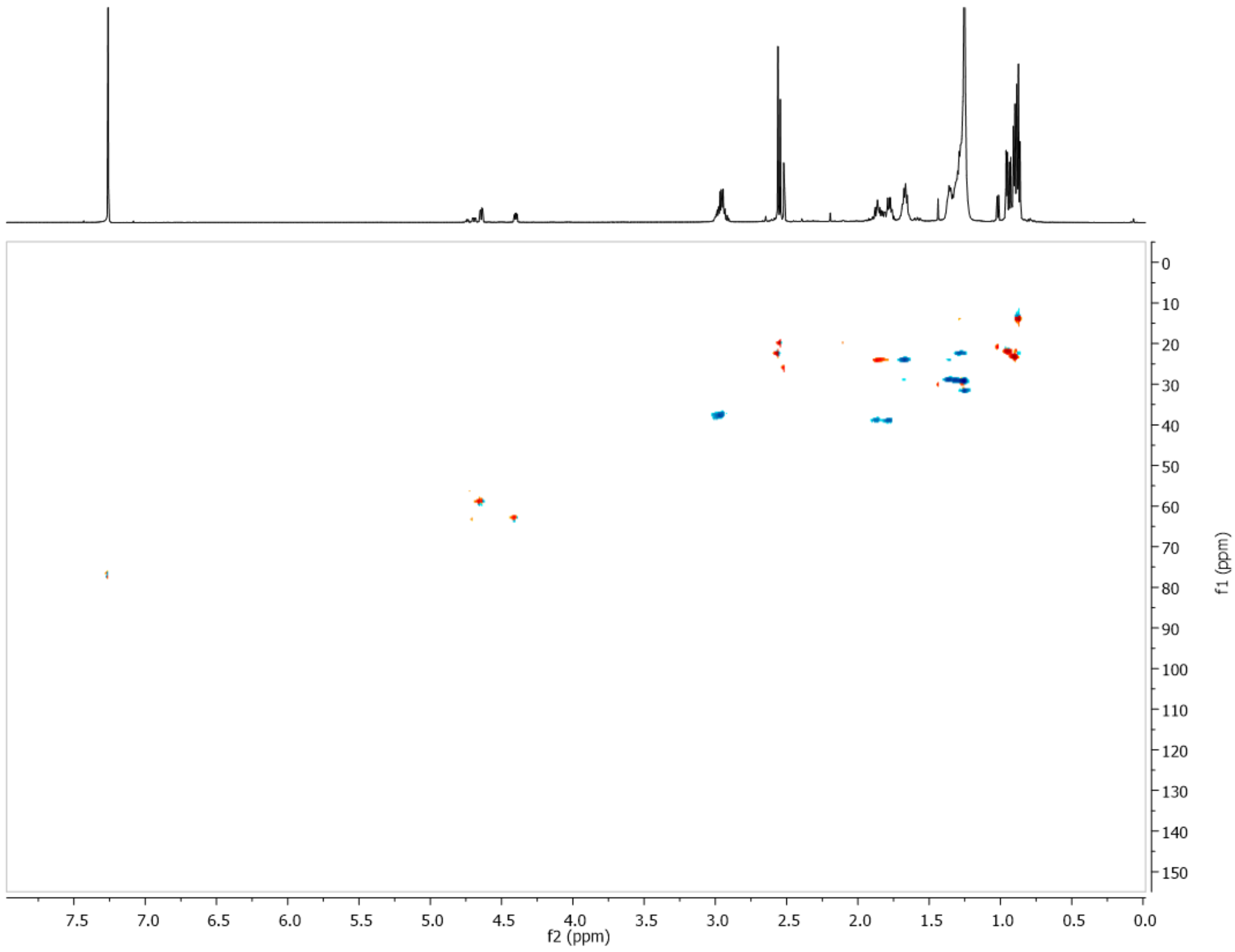

**Supplementary Figure 30.** HMBC spectrum of **5**.

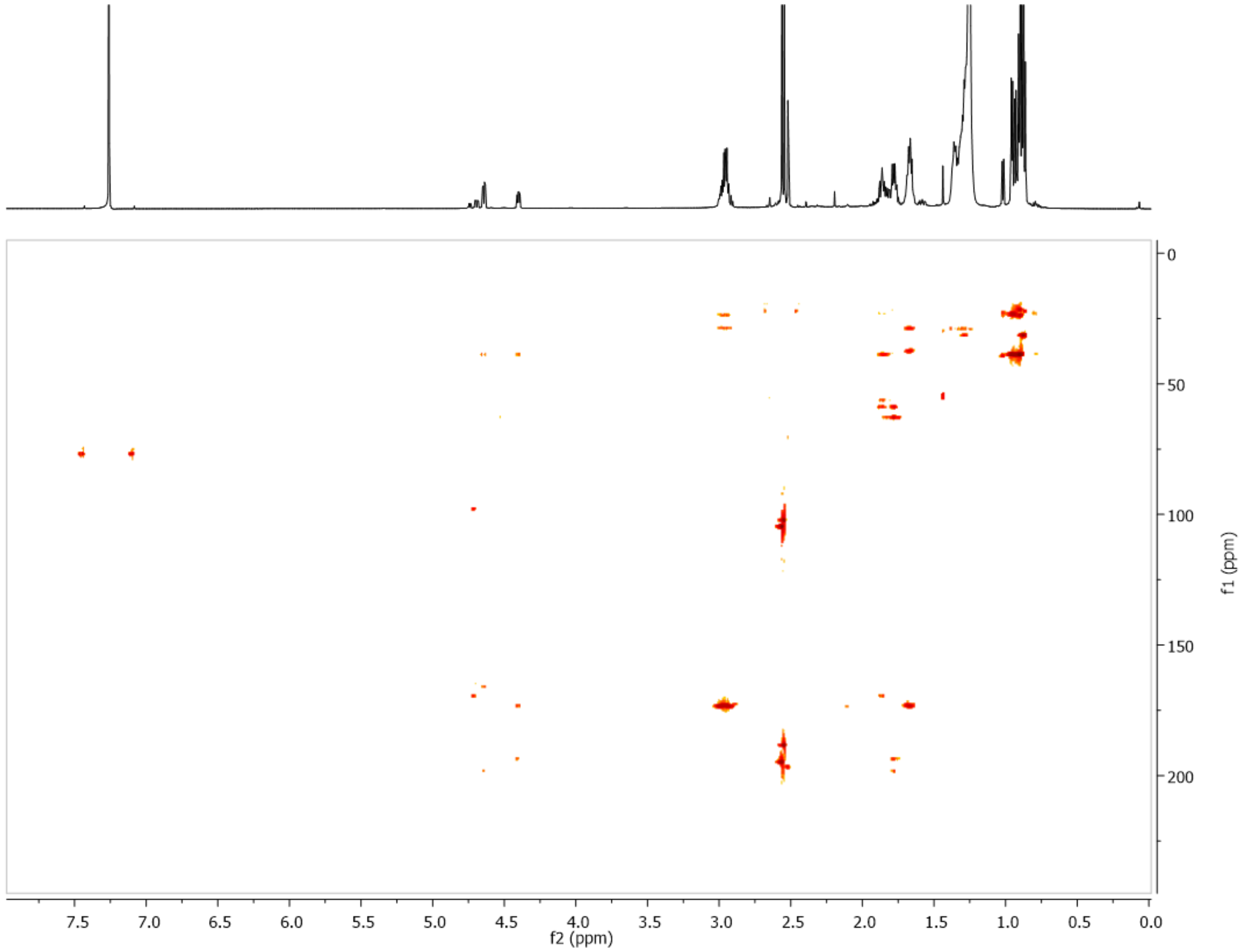

**Supplementary Figure 31.** Identification of free fatty acids from *S. mutans* B04Sm5.

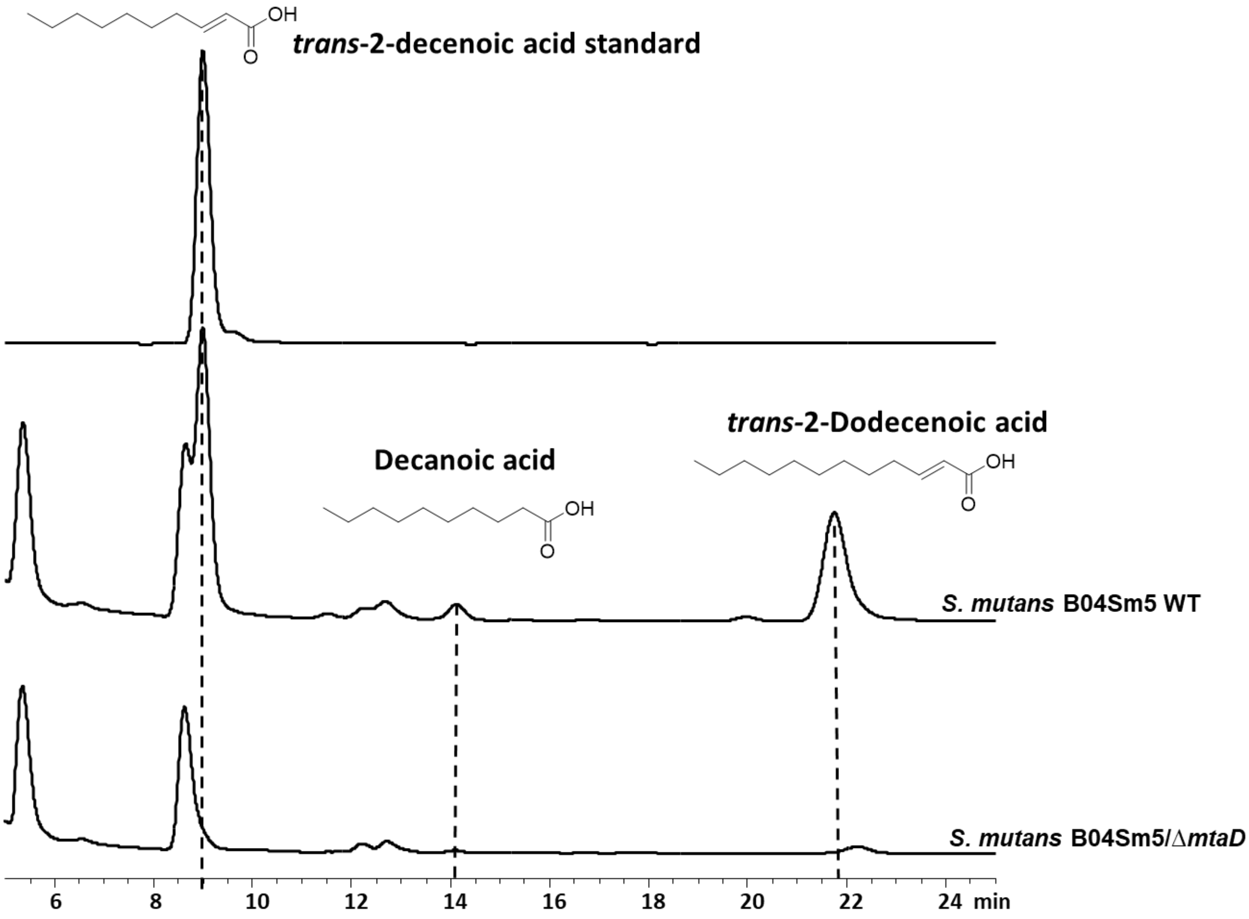

**Supplementary Figure 32.** MucF is predicted as a putitive membrane-bound protein.

### Phyre2

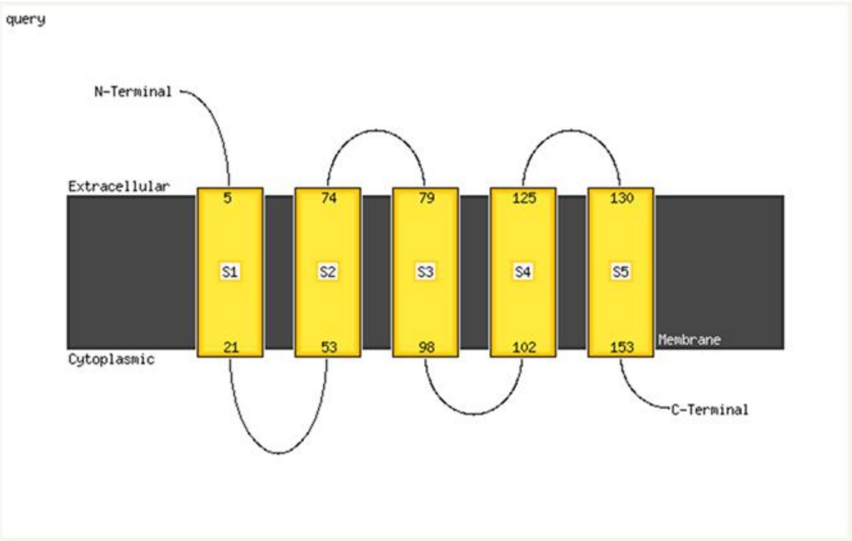

### Protter

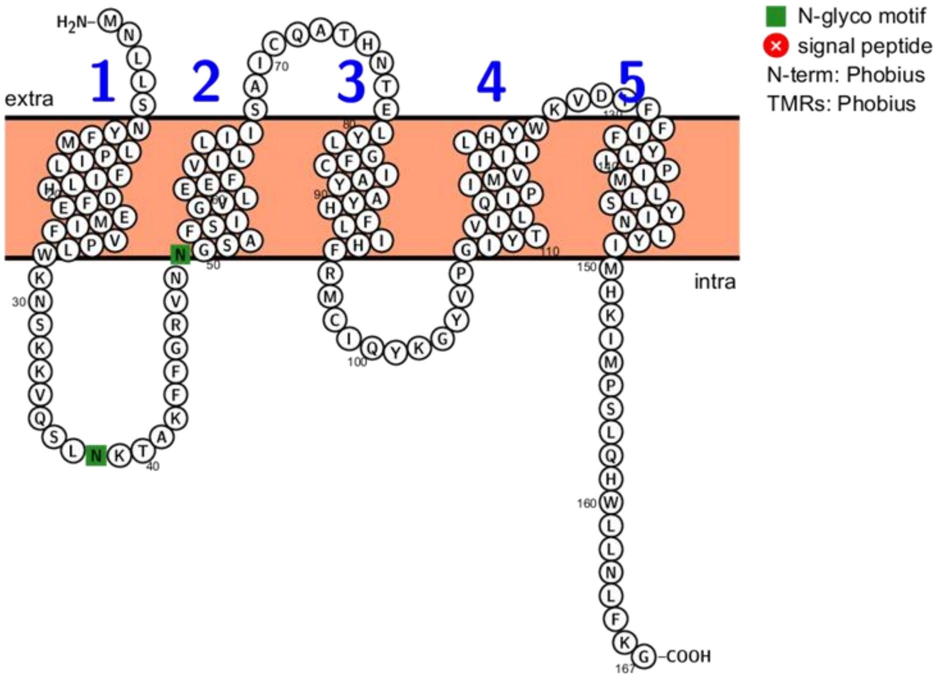

**Supplementary Figure 33.** Maximum likelihood (ML) tree of *S. mutans* B04Sm5 MucF (see asterics) and 498 homologous hypothetical proteins encoded by other bacterial genomes. Size of collpsed tree branches is proportional to the number of sequences within each clade. Bacterial taxa are labled at differnt levels of taxonomy ranging from species to phylum level (see color lables). CFB indicate the Cytophaga–Flavobacterium–Bacteroides phylum. Tree scale distance is presented as 0.1 (expected numbers of substitutions per site).

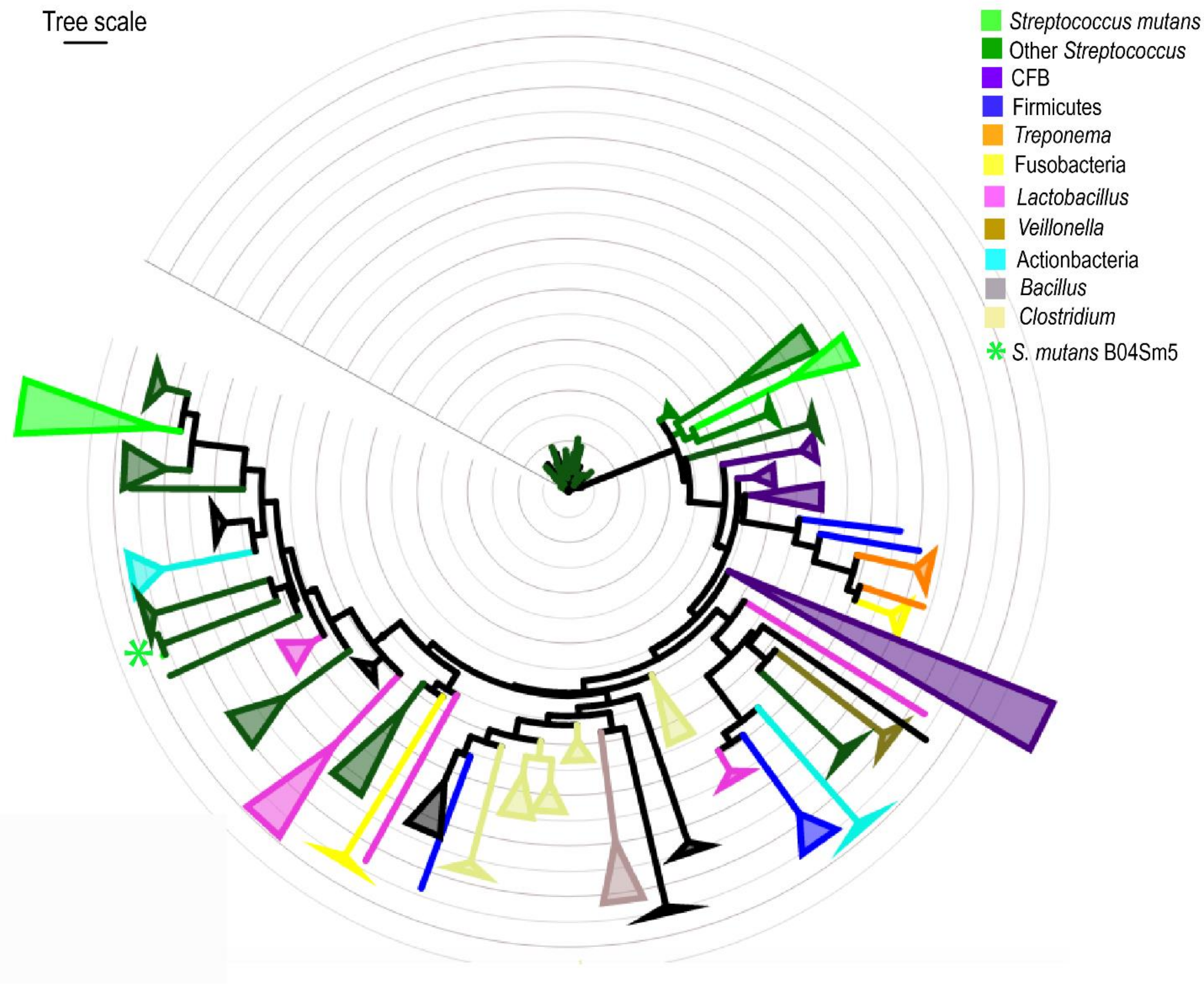
